# Nonlinear convergence boosts information coding in circuits with parallel outputs

**DOI:** 10.1101/811539

**Authors:** Gabrielle J. Gutierrez, Fred Rieke, Eric T. Shea-Brown

**Affiliations:** University of Washington, Department of Applied Mathematics; University of Washington, Department of Physiology and Biophysics

**Keywords:** Neural computation, Efficient Coding, Retina, Sensory Processing, Information Theory

## Abstract

Neural circuits are structured with layers of converging and diverging connectivity, and selectivity-inducing nonlinearities at neurons and synapses. These components have the potential to hamper an accurate encoding of the circuit inputs. Past computational studies have optimized the nonlinearities of single neurons, or connection weights in networks, to maximize encoded information, but have not grappled with the simultaneous impact of convergent circuit structure and nonlinear response functions for efficient coding. Our approach is to compare model circuits with different combinations of convergence, divergence, and nonlinear neurons to discover how interactions between these components affect coding efficiency. We find that a convergent circuit with divergent parallel pathways can encode more information with nonlinear subunits than with linear subunits, despite the compressive loss induced by the convergence and the nonlinearities when considered individually. These results show that the combination of selective nonlinearities and a convergent architecture - both elements that reduce information when acting separately - can promote efficient coding.

**Significance Statement:** Computation in neural circuits relies on a common set of motifs, including divergence of common inputs to parallel pathways, convergence of multiple inputs to a single neuron, and nonlinearities that select some signals over others. Convergence and circuit nonlinearities, considered individually, can lead to a loss of information about inputs. Past work has detailed how optimized nonlinearities and circuit weights can maximize information, but here, we show that incorporating non-invertible nonlinearities into a circuit with divergence and convergence, can enhance encoded information despite the suboptimality of these components individually. This study extends a broad literature on efficient coding to convergent circuits. Our results suggest that neural circuits may preserve more information using suboptimal components than one might expect.

---

Sensory systems, by necessity, compress a wealth of information gathered by receptors into the smaller amount of information needed to guide behavior. In many systems, this compression occurs via common circuit motifs - namely convergence of multiple inputs to a single neuron and divergence of inputs to multiple parallel pathways (1). Selective nonlinear circuit elements transform inputs, selecting some parts of the signal while discarding others. Here we investigate how these motifs work together to determine how much information is retained in compressive neural circuits.

These issues are highly relevant to signaling in the retina, because the bottleneck produced by the optic nerve ensures that considerable feedforward convergence occurs prior to the transmission of signals to central targets. This convergence reduces the dimension of signals as they traverse the retina. In total, signals from ∼100 million photoreceptors modulate the output of ∼1 million ganglion cells (2, 3). If the dynamic range of the ganglion cell is not sufficiently expanded beyond that of the photoreceptors and bipolar cells, this convergent circuit architecture could lead to a compression of input signals in which some information or stimulus resolution is lost - resulting in ambiguously encoded stimuli. It is estimated that the population of ganglion cells collectively transmits approximately 10^6^ bits of information (3–5) and that this is much less than the amount of information available to the photoreceptors (2). However, not much is known about how neuron properties interact with a convergent circuit structure to drive or mitigate a loss of information.

Receptive field subunits are a key feature of the retina’s convergent circuitry. Multiple bipolar cells converge onto a single ganglion cell - forming functional subunits within the receptive field of the ganglion cell (6, 7). Ganglion cell responses can often be modeled as a linear sum of a population of nonlinear subunits. These subunit models have been used to investigate center-surround interactions (8–12) and to explain the nonlinear integration of signals across space (7, 10, 13–15).

While it is clear that subunits have the potential to compress inputs, it is not known whether this architecture subserves an efficient code where inputs are encoded with minimal ambiguity. For decades, information theory (16, 17) has been used to quantify the amount of information that neurons encode (3, 5, 18–27). The efficient coding hypothesis proposes that the distribution of neural responses should be one that is maximally informative about the inputs (21, 22, 28). Take the example of a stimulus variable, such as luminance, where the brightness level is encoded by the number of spikes in the response. An input/output mapping in which most of the possible luminance levels are encoded by the same response (i.e. the same number of spikes or firing rate) makes many bright and dim inputs ambiguous and provides very little information. Information can be maximized at the level of a single neuron by distributing the responses such that they optimally disambiguate inputs (23). A nonlinear response function optimized for the distribution of inputs can make the most of the neuron’s dynamic range. Adaptive rescaling of the response nonlinearity to changes in the input statistics can maintain maximal information in the output (29–31). Alternatively, information can be maximized by optimizing connection weights in the circuit, perhaps in combination with optimizing the nonlinearities (19, 32, 33). These past works, however, have not made explicit how the set of motifs found in most neural circuits, and in the retina in particular, combine to collectively influence coding efficiency.

Our contribution here is to dissect a canonical neural circuit in silico, and to investigate how much each of its components contribute to or detract from the information encoded by the circuit about stimuli. These circuit components, considered individually, have the potential to discard information. We begin with the simplest motif of converging inputs to single neurons, and analyze the role of rectifying nonlinear subunits applied to each of these multiple inputs. We then add a diverging motif which splits the response into two opposing pathways. We find that rectifying nonlinear subunits mitigate the loss of information from convergence when compared to circuits with linear subunits. This is despite the fact that the rectifying nonlinear subunits, considered in isolation, lead to a loss of information. Moreover, this ability of nonlinear subunits to retain information stems from a reformatting of the inputs to encode distinct stimulus features compared with their linear counterparts. Our study contributes to a better understanding of how biologically-inspired circuit structures and neuron properties combine to impact coding efficiency in neural circuits.

## Results

We start by quantifying the effect of common circuit motifs, alone and in combination, on coding efficiency. We then explore, geometrically, how nonlinear subunits shape the response distribution to gain intuition as to how they can lead circuits to retain more information. Finally, we explore the implications of nonlinear subunits for encoding stimulus properties. To emphasize the geometrical characterization of the encoding, we use an abstract circuit model without temporal dynamics.

### Common circuit components are lossy or inefficient

Our goal is to understand how the combination of divergence of inputs and convergence of nonlinear subunits impacts the retina’s ability to efficiently encode spatial inputs. We are particularly interested in the impact of selective nonlinearities on efficient coding. We use Shannon’s information to describe the maximum amount of information that a distribution of responses could contain about its inputs (16, 34). We consider deterministic circuits in which the mutual information between the stimulus and response reduces to the entropy of the response. Specifically, we use discrete entropy to compare the information content of continuous distributions of responses generated by different model circuits. We also confirm our results by computing the mutual information of noisy circuit responses (see SI Appendix). The parameters of the discretization were chosen so that the difference between the area under the discretized distribution and its continuous counterpart was minimized for a range of distinct distributions (see Methods).

Many neural circuits are organized in layers of converging and diverging neurons and connections. In the retina (Fig. 1A), this produces a compression and “re-formatting” of a highdimensional visual input into a lower dimensional neural code that can be interpreted by the brain. In addition, nonlinear responses abound in the neurons that compose these layers. These mechanisms may complicate the ability of the circuit to retain information. For example, two converging inputs can result in ambiguities. With linear convergence, the ability to distinguish the stimulus combinations that sum to the same value is lost and hence this is a form of lossy compression (Fig. 1B). The entropy of the full two-input stimulus (Fig. 1B, top) is 14.68 bits - meaning that a given point in the stimulus space provides 14.68 bits of information about the identity of the stimulus (given our choice of bin size, see Methods). The entropy of the convergent response is smaller (7.87 bits; Fig. 1B, bottom), thus indicating ambiguity in the stimulus identity.

**Fig. 1.**
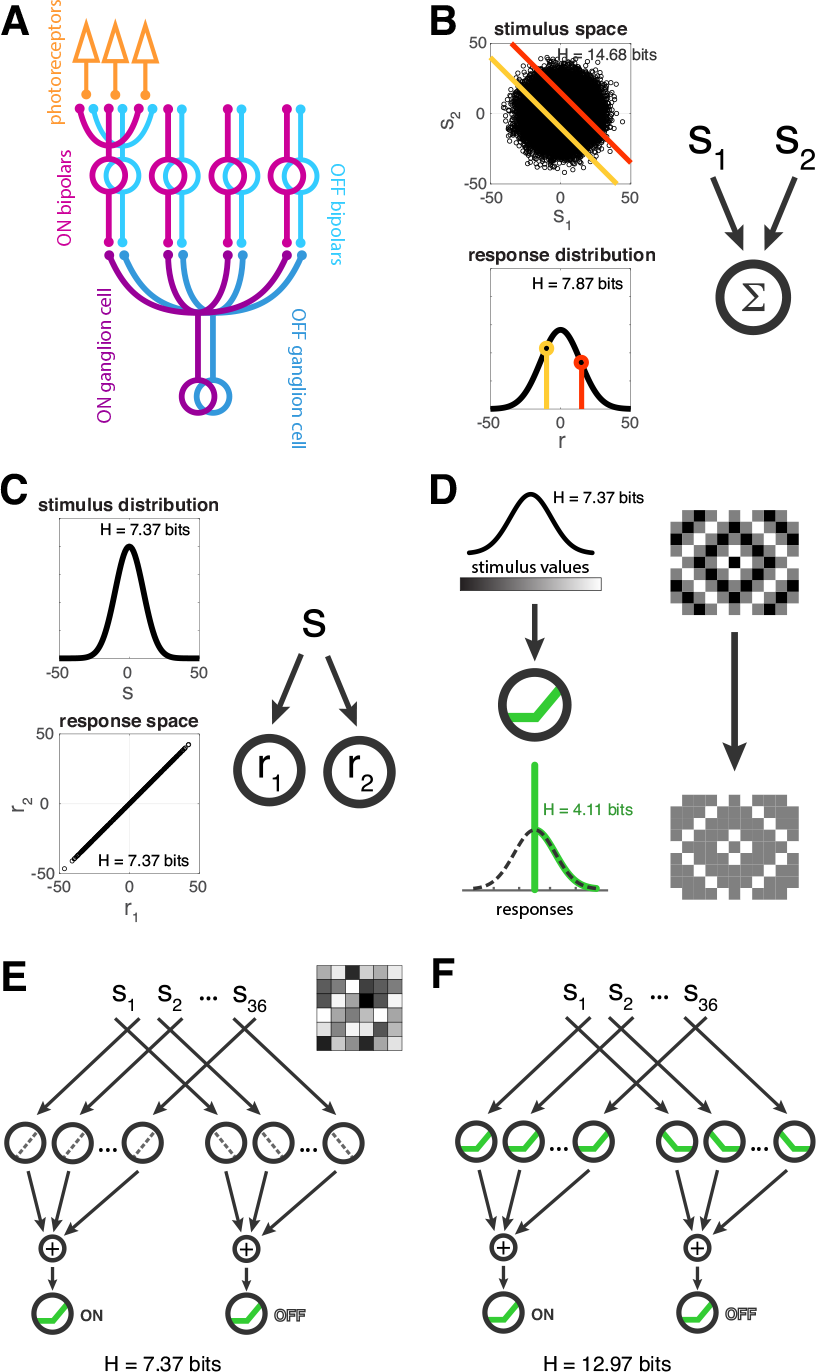
Neural circuits are composed of inherently lossy components. (A) Schematic of retina circuit with its convergent and divergent structure. (B) Converging two inputs results in ambiguities. A 2-input stimulus space is reduced to a single output response space in which one response (bottom: yellow and orange points) represents all stimuli along an isoline (top: yellow and orange lines) where *s*_1_ + *s*_2_ = constant. All entropy values shown are based on a discrete entropy computation (see Methods). (C) Diverging a signal to two outputs can produce redundancies. (D) Nonlinear transformation of a gaussian distributed stimulus input with a ReLU (rectified linear unit) can distort the distribution, producing a compressed response in which some portion of the stimulus information is discarded. (E-F) Convergent, divergent circuits with (E) linear subunits, or (F) nonlinear subunits. Subunit responses are weighted by 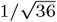 Example stimulus image is shown.

Diverging motifs are another common neural circuit construction. In the example shown in Figure 1C, the divergent responses are identical and the entropy of the 2-dimensional response space (H = 7.37 bits) is the same as the entropy of the 1-dimensional stimulus distribution shown in the top plot (H = 7.37 bits). This demonstrates that divergence of an input into two neurons may produce an inefficient neural architecture by producing redundant or correlated signals.

Nonlinearities are abundant in neural circuits, and firing rates generally have a nonlinear relationship to inputs. On a more granular level, synaptic and spike generation mechanisms are often nonlinear and can be approximated by thresholded functions. The rectified linear nonlinearity is a tractable representation that captures key features of neural nonlinearities, including the selectivity for some inputs over others. The subunits in our model most closely represent bipolar cells, and we interpret the subunit nonlinearities as the relationship between input and excitatory synaptic output. The output units in our model most closely represent ganglion cells, and we interpret the output nonlinearities as occurring in spike generation.

Similarly to convergence, nonlinear transformations can lead to loss of information by introducing ambiguities. Take the example of a rectified-linear transformation that is thresholded at zero and is therefore selective for positive inputs (Fig. 1D). It is a non-invertible nonlinearity where half of the stimulus distribution is encoded faithfully and half is mapped to an output of 0 by the thresholded response. Therefore, this nonlinearity induces lossy compression: the information that would distinguish these thresholded stimuli has been irretrievably discarded. Correspondingly, the entropy of the rectifying nonlinear response (H = 4.11 bits) is around half of that for the stimulus distribution (H = 7.37 bits).

Each of the common circuit motifs described above is inefficient or discards information when considered in isolation (Figs. 1A-D). How much information can a neural circuit with all of these components retain? We constructed a model circuit that compresses a high-dimensional spatial input into a low-dimensional output. It has an N-dimensional input structure that diverges along two pathways, an ON and an OFF pathway, each culminating in a single output neuron. The inputs to each output neuron come from a layer of subunits - the building blocks for the receptive field structure of the output neuron. Each subunit receives input from one of the N stimulus inputs that compose a stimulus image, and each stimulus input is independently drawn from a gaussian distribution. Within each pathway, the normalized subunit responses linearly sum at the output neuron and are then rectified.

The ON and OFF output responses lie in a 2-dimensional space, and form a low-dimensional representation of the N-dimensional input. We compute the entropy of the 2-dimensional output response after showing many stimulus samples to the circuit. In our study, circuits have rectifying output neurons which model the rectifying responses of ganglion cells. We wanted to know whether the subunits - which represent non-spiking bipolar cells - reduce or enhance the information encoded by the circuit when their responses are also rectifying compared to when subunit responses are linear. For a 36-dimensional input space, the circuit with linear subunits (LSC: linear subunits circuit) has 7.37 bits of entropy (Fig. 1E), while the circuit with nonlinear subunits (NSC: nonlinear subunits circuit) has 12.87 bits of entropy (Fig. 1F). The greater entropy of the NSC is counterintuitive because the nonlinear neurons considered in isolation lead to a loss of information (Fig. 1D).

This unexpected result motivated us to consider how each circuit component interacts with the others to determine the encoded information. Our claim is that nonlinear subunits, together with nonlinear output neurons, retain more information than linear subunits together with nonlinear output neurons. This necessarily differs from a claim that the NSC produces more information than what is available in the stimulus, as no processing operation can increase the information content of an input signal (Data Processing Inequality, 17). Neither circuit in Figure 1E,F retains the full amount of information in the 36-dimensional input signal which has a much higher entropy (H = 265.28 bits, see Methods) than the 2-dimensional outputs produced by either circuit. The convergence of the inputs necessarily limits the information in the output (17). To illustrate, the population of ON linear subunits contains the same amount of information as the stimulus; however, that information will be reduced as soon as the subunits are summed. The rectification that follows the summation will further reduce the encoded information. In contrast, at the level of the population of nonlinear subunits, the encoded information will be reduced early on by the rectifying subunits. The same summation and output rectification follows, and the net result is that there is less information lost in the final output. This advantage is due to nonlinear processing at the subunit level.

Our study concerns the reformatting of stimulus information by nonlinear subunits. We chose nonlinearities that are inherently selective for parts of the stimulus inputs (i.e. ON, OFF rectification) as a generic model for the selectivities in bipolar cells. As discussed above, such selective nonlinearities discard information at the single neuron level. Rather than optimizing circuit weights or input biases to maximize information, our goal is to explore the contribution of generic, fixed nonlinearities that operate independently on signals in each subunit within a parallel circuit. We next investigate how convergence interacts with these subunit nonlinearities.

### Lossy nonlinear subunits benefit from convergence

To understand the joint impact of nonlinear subunits and convergent connectivity on encoded information, we examined circuit configurations with a single pathway, i.e. without divergence (Fig. 2). Pathways with two subunits permit visualization of the input and response spaces. Stimuli that sum to the same value (example highlighted with dark purple in the top plot of Fig. 2A) elicit the same response in the circuit pathway with linear subunits because the subunits do not transform the inputs (Fig. 2A, left, 3rd and 4th rows). The nonlinear subunits transform the stimulus space such that all points are compressed into a single quadrant (Fig. 2A right, 2nd row). Summing the nonlinear subunits (Fig. 2A, right, 3rd row) allows the potentially ambiguous stimuli to have a more distributed representation in the output response - meaning that they are represented more distinctly by the nonlinear subunits pathway than the pathway with linear subunits.

**Fig. 2.**
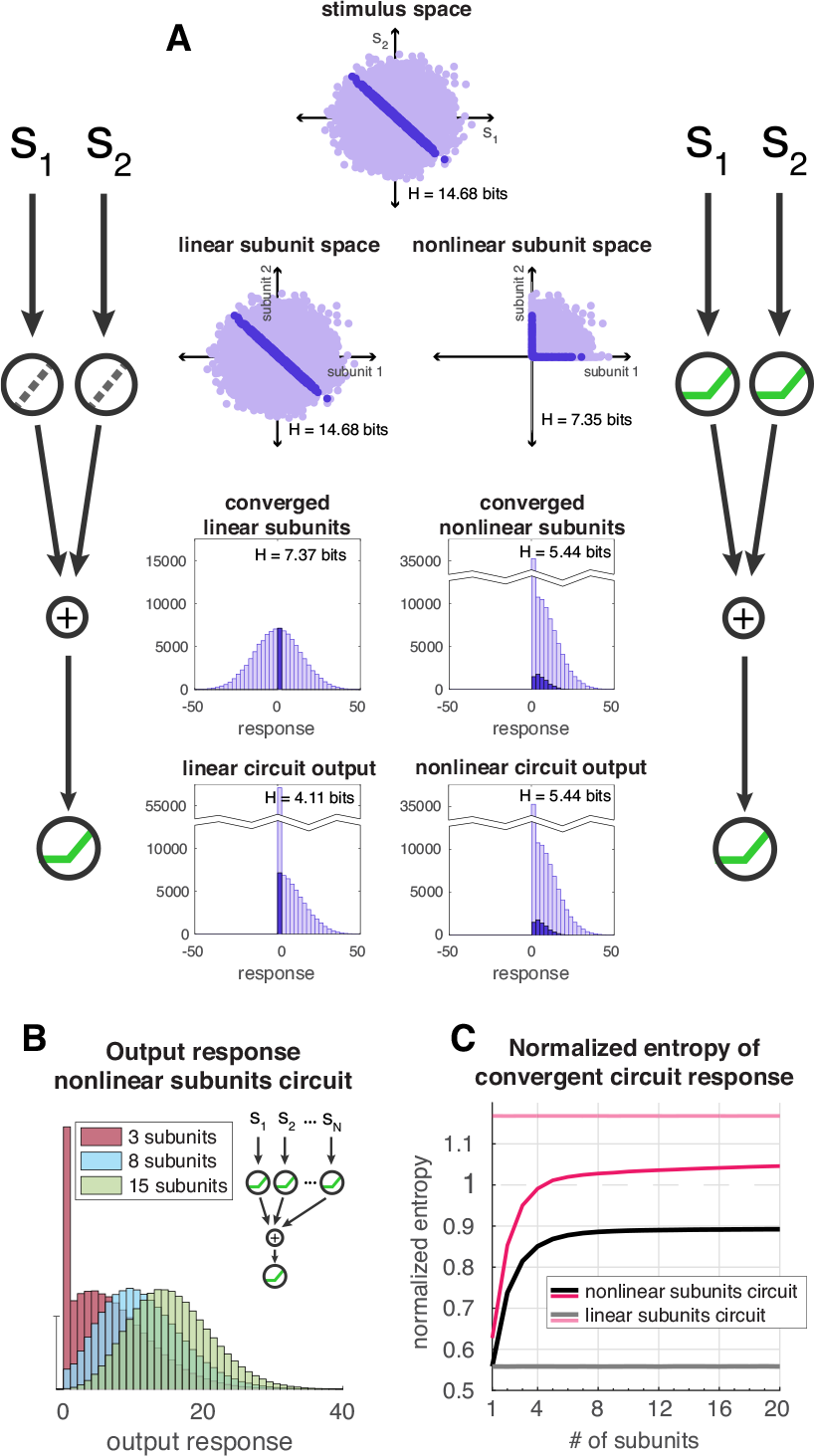
(A) The encoding of the stimulus space (top) within each layer of a 2-subunit convergent circuit configuration without divergence. Subunits (2nd row); summed subunits response distribution (3rd row); nonlinear output response distribution (4th row). Left, linear subunits circuit; right, nonlinear subunits circuit. The output nonlinearity does not have an additional effect on the summed nonlinear subunits without noise. (B) Histograms of the output response for the NSC are shown for configurations with 3, 8, and 15 subunits. The subunit responses are normalized so that each subunit is weighted by 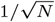 where *N* is the number of subunits. The inputs [*s*1, *s*2, …, *sN*] are independently drawn from a gaussian distribution (see Methods). (C) Normalized entropy of output response as a function of number of convergent subunits where subunits are normalized as in B and the circuit entropy is normalized by the entropy of the summed linear subunits (see Methods). Black curve, NSC; gray, LSC; dark pink, nonlinear subunits with optimized sigmoidal output nonlinearity; light pink, linear subunits with optimized sigmoidal output nonlinearity. Standard deviation of entropy over 10 runs for each configuration is on the order of between 10^*–*4^ and 10^*–*2^ bits.

For a configuration with a single subunit, the LSC and NSC would have identical output responses so long as there remained an output nonlinearity. The 2-subunit circuit (Figure 2A) showed improved information transmission with nonlinear subunits over linear subunits, and this prompted us to ask whether there would be a continued improvement with additional nonlinear subunits. We computed the entropy of the output responses for the linear and nonlinear subunit configurations that converge to a single output for a range of subunit quantities (Fig. 2B,C; also see SI Appendix, Fig. S1A). With increasing numbers of subunits, more subunit responses are converged into the output response. To observe a relative change in entropy as the number of subunits is increased, the subunits were normalized; and to observe the dependence of this effect on the nonlinearities, the output response entropy was normalized by the entropy of the summed linear subunits (see Methods).

The distribution of output responses for the nonlinear subunits pathway qualitatively changes with the number of subunits (Fig. 2B). With few subunits, the output response distribution resembles the truncated gaussian seen for the rectified output response in Figures 1D and 2A. With increasing numbers of subunits, the output response distribution approximates a gaussian (due to the central limit theorem) with a mean that shifts towards more positive values (Fig. 2B; also see SI Appendix, Fig. S2).

The entropy for the nonlinear subunits pathway increases with increasing subunit dimension (Fig. 2C, black line). It saturates near a normalized value of 0.9, before ever reaching the entropy of the converged linear subunits (where normalized entropy is 1); thus, although increasing convergence improves the information retention of nonlinear subunits, the entropy of the converged nonlinear subunits is apparently bounded by the entropy of the converged linear subunits. The nonlinear subunits only encode positive inputs whereas the linear subunits encode positive and negative inputs. However, when the summation of the linear subunits is followed by a nonlinear rectification at the output, the response entropy is reduced (H = 4.11 bits, Fig. 2C, grey line, normalized H = 0.56) and does not increase beyond that regardless of the number of convergent subunits.

The output nonlinearity reduces the entropy of the LSC whereas in the NSC the output nonlinearity does not impact the entropy of the summed nonlinear subunits since the responses have already been rectified. The summed linear subunits produce a gaussian distribution, and the summed nonlinear subunits approach a gaussian distribution as greater numbers of subunits are converged. The entropy of either circuit could be maximized by replacing the output nonlinearity with a sigmoidal nonlinearity that is the cumulative gaussian of the summed subunits distribution bounded by the maximum and minimum values of that distribution (23, see Methods). Doing so benefits the linear subunits motif more than the rectified subunits motif (compare dark and light pink curves, Fig. 2C; and in SI Appendix, Fig. S1A) because the variance of the full distribution of summed linear subunits is greater than that for the distribution of summed nonlinear subunits.

Figure 2 illustrates how the placement of the rectified nonlinearity within the circuit impacts the entropy of the response. When the nonlinearity is placed within the subunits, less information is lost than when the nonlinearity is shifted further down in the circuit after the summation of linear subunits. These results continue to hold for the mutual information between the output response and the stimulus when noise is added after the subunit summation (SI Appendix, Fig. S1). We wondered whether this effect of nonlinear convergence was sufficient to explain why the divergent NSC in Figure 1F has higher entropy than the divergent LSC (Fig. 1E). We next explore the impact of divergence on information coding with nonlinear subunits.

### Divergent circuit structure leverages selectivity of nonlinear subunits

To understand the combined impact of divergence, convergence, and nonlinearities, we present a geometrical exploration of the transformations that take place in the different layers of the circuit with either linear or nonlinear subunits. Our demonstration uses circuits with two input dimensions to facilitate visualization of the stimulus and subunit spaces (Fig. 3).

**Fig. 3.**
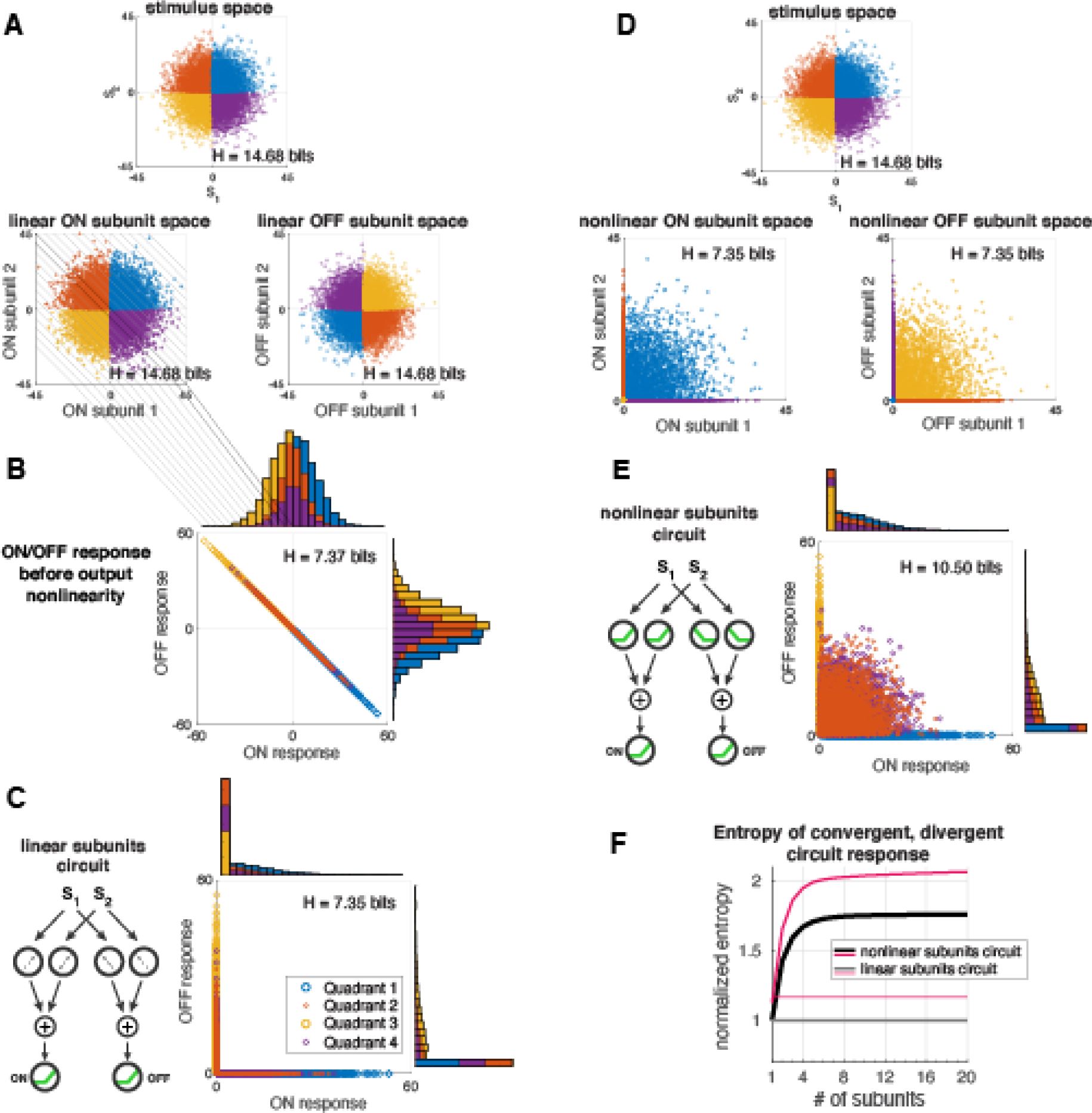
Visualization of stimulus and response mappings at each level of a convergent, divergent circuit with two inputs, two subunits for each pathway (ON and OFF pathways), and a nonlinear output neuron for each pathway. The points in all subsequent plots are color-coded by the stimulus quadrant from which they originate. (A) The stimulus space (top) has color-coded quadrants. The 2-input stimulus space maps onto a 2D linear subunit space for each pathway (second row, left: ON; right: OFF). The subunit spaces are shown before subunit normalization. (B) The response space is shown for the linear sum of subunits before the output nonlinearity is applied and (C) after the nonlinear output response. (D) The 2-input stimulus space (top) maps onto a 2D **nonlinear** subunit space for each pathway (second row, left: ON; right: OFF). (E) The output response space for the NSC. Note that the output response before the output nonlinearity is applied (not shown) is identical to the output response after the output nonlinearity is applied for the circuit with nonlinear subunits. (F) Normalized entropy of the output response for convergent, divergent circuits with increasing input and subunit dimension (subunit responses are normalized as before). The circuit entropy is normalized by the entropy of the summed linear subunits. Gray, LSC in C; black, NSC in E; light pink, LSC with optimal sigmoidal output nonlinearity; dark pink, NSC with optimal sigmoidal output nonlinearity.

To determine the optimal nonlinear thresholds, we swept through a range of thresholds for ON and OFF subunits in a divergent, convergent circuit with two inputs and computed the response entropy for each combination of threshold values.

Very low thresholds approximate linear functions while high thresholds are extremely rectifying. We found that the optimal combination of ON and OFF subunit thresholds meet at zero (SI Appendix, Fig. S3). These zero-crossing nonlinearities are used for all other figures in the main text. Furthermore, when output responses are considered abstractly as static firing rates, this position of thresholds produces low mean output responses that are comparable to those from the most rectified subunits (SI Appendix, Fig. S2 and S4).

As before, the linear ON subunit space (Fig. 3A, 2nd row, left) is identical to the stimulus space (Fig. 3A, top) because no transformation or compression has taken place through the linear subunits. The OFF subunits receive a negative copy of the same stimulus that the ON subunits receive which reflects the stimuli about the diagonal (Fig. 3A, 2nd row, right). When the linear subunits converge within their respective pathways, the ON and OFF responses are compressed onto a diagonal line because they are anti-correlated (Fig. 3B). This emphasizes the fact that the ON and OFF linear subunits do not have stimulus selectivities in the strictest sense. When the output nonlinearities are applied, this linear manifold is folded into an L-shape (Fig. 3C).

The entropy for the output response of the LSC with diverging pathways (H = 7.35 bits) is higher than it was with just a single pathway (H = 4.11 bits, Fig. 2A). However, it is only increased enough to nearly match the entropy of a single pathway response without any nonlinearities in either the subunits or the output (H = 7.37 bits). In other words, the OFF pathway in the LSC with output nonlinearities (Fig. 3C) encodes the information discarded by the output nonlinearity in the ON pathway, but it does not enable the divergent LSC in Figure 3C to do any better than the convergence of only ON linear subunits (Fig. 2A). This is because the linear subunits do not select for anything specific and nothing is lost to selectivity; instead the loss of entropy (relative to the entropy of the stimuli) occurs from convergence. Furthermore, when the convergence of the ON linear subunits is followed by a nonlinearity, only the positive-summing stimuli are selected. A divergent OFF pathway selects the negative-summing stimuli that the ON pathway discards. Visually, one can see that nothing is lost by folding the linear response space into an L. The divergent LSC recovers what is lost by the output nonlinearities, but not what is lost by convergence.

Unlike the linear subunits, the stimulus undergoes a transformation within the nonlinear subunits layer (Fig. 3D), producing a complimentary compression for the ON and OFF pathways. When these subunits converge in their respective pathways (Fig. 3E), the output response has some similarities to that for the LSC (Fig. 3C). The L-shaped manifold is still present, but the points representing the stimulus inputs with mixed sign have been projected off it. By virtue of having these points leave the manifold and fill out the response space, entropy is increased. In fact, as more nonlinear subunits converge in a divergent circuit, a greater portion of points are projected off the manifold along the axes, and as a result the entropy continues to increase until saturation (Fig. 3F, black curve). These results continue to hold for the mutual information between the output response and the stimulus when independent noise is added to the subunit summation in each pathway (SI Appendix, Fig. S1B).

The NSC does nothing to save the dually positive (blue quadrant) or dually negative (yellow quadrant) stimuli from information loss by convergence. Those are ultimately encoded in the same way as by the LSC. In fact, the circuit entropy is less sensitive to the subunit thresholds when the stimuli corresponding to different subunits are correlated (i.e. between *s*_1_ and *s*_2_) than when the stimuli are anti-correlated (SI Appendix, Fig. S5). The advantage conferred by the divergent nonlinear subunits is to preserve the variance among the mixed sign stimuli, not only within a single pathway, but also across ON and OFF pathways (this is why adding a bias to the summed linear subunits to evade the output nonlinearity will not match or surpass the entropy of the NSC). As the stimulus dimension is increased, the mixed sign stimuli make up a larger and larger proportion of all stimuli, resulting in the increasing advantage of the NSC and its saturation.

To show that the nonlinear subunits themselves confer a unique advantage, we once again replace the output nonlinearities with optimal sigmoidal nonlinearities that are the cumulative gaussian of the summed subunit distribution. The entropy of the LSC is increased (Fig. 3F, light pink), however, it is not increased beyond the entropy of the NSC with (Fig. 3F, dark pink) or without (Fig. 3F, black) optimal output nonlinearities. This demonstrates that the entropy of the convergent, divergent circuit can be increased beyond an optimization of the output nonlinearities by implementing selective nonlinear subunits.

The rectified output nonlinearities have the effect of decorrelating the ON and OFF output responses in the LSC, while for the NSC, it is the nonlinear subunits themselves that decorrelate the output responses (correlation coefficients: linear response = -1, Fig. 3B; LSC = -0.4670, Fig. 3C; NSC = -0.4669, Fig. 3E). Indeed, although the output nonlinearity decorrelates the ON and OFF outputs of the LSC, this decorrelation does not produce any gains in entropy relative to the LSC before output nonlinearities are applied. Furthermore, the ON and OFF responses of the NSC are as decorrelated as for the LSC, but unlike the LSC, it experiences an entropy gain over the converged linear subunits alone. Complementing the geometrical explanations above, SI Appendix II presents an analytic argument for why the NSC has greater entropy than the LSC using the fact that the summed subunit distributions in both circuits are gaussian in the limit of large N subunits.

The additional entropy conferred by divergence for the NSC is due to *how* the nonlinear subunits decorrelate the ON and OFF pathways, and not merely the fact that those pathways have been decorrelated. It is this subunit processing step that pulls responses off the linear manifold in the output response space leading to an increase in response entropy. The space of the responses in the linear case can be expanded by manipulating the linear subunit weights; however, we find that no rotation of the linear subunit weights can cause the entropy of the LSC to surpass that of the NSC (SI Appendix, Figs. S6 and S7). Furthermore, decorrelating the nonlinear subunit weights confers limited benefit relative to decorrelating the linear subunit weights (SI Appendix, Fig. S8).

To determine whether the increase in entropy for the NSC is due to a “synergistic” effect whereby the ON and OFF output responses convey more information together than the sum of the information that each output contains individually (3, 35, 36), we computed the synergy (*syn*(*R*_1_, *R*_2_) = *I*(*S*; *R*_1_, *R*_2_) *– I*(*S*; *R*_1_) *– I*(*S*; *R*_2_)) for the different circuit configurations and for a range of subunit quantities (SI Appendix, Fig. S9). Positive values of this metric indicate synergy while negative values indicate redundancy. None of the circuits have synergy; however, the NSC has less redundancy than the LSC.

Increased response entropy could reflect an increased precision in encoding the same stimulus features or the encoding of new stimulus features. We next explore how the processing of mixed sign stimuli by nonlinear subunits creates sensitivity to stimulus features that are not encoded with linear subunits.

### Nonlinear subunits circuit encodes both mean and contrast information

To determine whether the boosted entropy of the NSC accompanies an encoding of additional stimulus features, we visualized the stimulus and response spaces for the linear and nonlinear circuit configurations. The stimulus inputs are assumed to represent luminance values and the distributions are the same as before. We chose two basic features of visual stimuli to investigate: mean luminance and contrast. In Figure 4A, the stimulus space is color-coded by bands of mean luminance levels. In the response spaces for the LSC and NSC a banded structure is preserved (Fig. 4A), indicating that there is a separation of the mean luminance levels within the response spaces for both circuits.

**Fig. 4.**
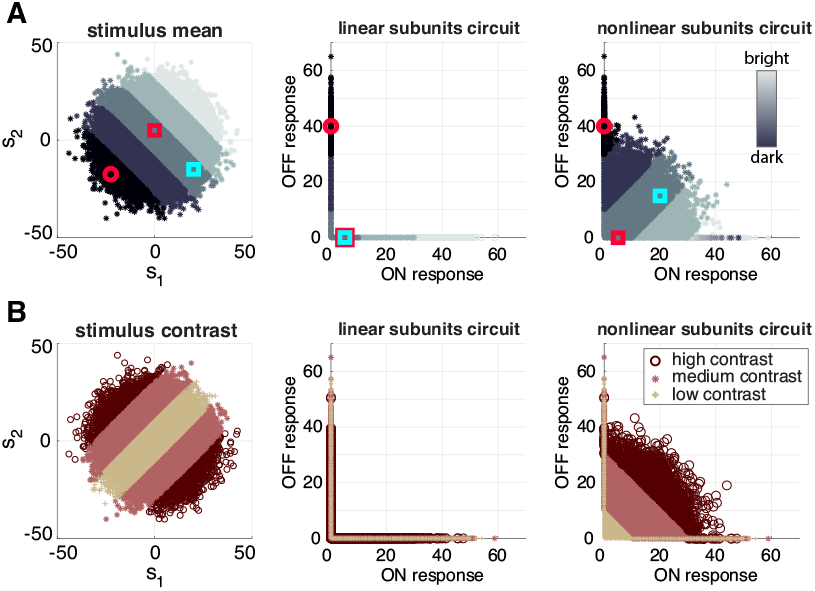
Mean and contrast encoding of convergent, divergent circuits from Figure 3 (2 inputs, ON and OFF outputs). (A) Visualization of the stimulus mean and output response spaces. For example, the bright mean stimulus band contains the 2-input image samples that have the highest mean luminance. The red square is an arbitrary reference point. In the stimulus space, the cyan square has the same mean luminance as the red square but a different contrast, while the red circle has the same contrast as the red square but a different mean luminance. (B) Visualization of the stimulus contrast and output response spaces. The high contrast stimulus bands contain 2input image samples that have high contrast, whereas the low contrast band contains 2-input image samples where the input luminance is more correlated.

Contrast is encoded differently the two circuits (Fig. 4B). The stimulus space in Figure 4B (left) is color-coded for three contrast levels. The highest contrast areas of the space are in the mixed sign quadrants. The representations for low, medium, and high contrast stimuli overlap each other in the output response space of the LSC (Fig. 4B, middle). However, there is separation of these contrast levels in the output response space of the NSC (Fig. 4B, right). As the number of inputs increase, so too does the proportion of mixed sign inputs, giving the NSC a continued advantage in encoding contrast over the LSC as more subunits are converged. This is reinforced by the result that the NSC is more sensitive to anti-correlated stimuli than the LSC (SI Appendix, Fig. S5, right panel). Thus, the NSC encodes both mean and contrast information whereas the LSC only encodes mean luminance.

## Discussion

In a circuit like the retina, inputs diverge to distinct cell types while neurons receive converging inputs from many presynaptic neurons. This combination of divergence and convergence reorganizes and compresses visual inputs. To determine the impact of these common circuit properties on information encoding, we built a circuit model and compared the response entropies of linear and nonlinear subunit configurations. Divergence, convergence, and non-invertible nonlinear signal transformations each have a negative impact on efficiency, or information, individually. However, when arranged together they can mitigate the loss of information that is imposed by the reduction in dimension from inputs to outputs.

The advance made by our study is to demonstrate that rectified nonlinearities can increase the response entropy in a circuit with convergence and divergence, not merely by decorrelating inputs, but by re-coding them. We predict that the information encoded by neurons is maximized by circuit mechanisms that exploit such nonlinearities before inputs converge. This complements known mechanisms, such as adaptation and response equalization, that enhance coding efficiency by providing a good match between input stimuli and responses at the level of the output neuron (23, 30–32, 37).

### Transforming convergent inputs enhances circuit efficiency

For a single neuron receiving a single input with a known distribution, classical and influential studies prescribe how transmitted information can be maximized by matching the response function to that distribution (23, 30, 31). We consider a complementary question here: when there are multiple inputs converging to a neuron, how should those inputs be transformed to maximize the information that a neuron – or that multiple neurons within a divergent output population – can transmit?

The weights with which inputs are combined is often a key factor in the information encoded by a circuit and this issue has been studied extensively (19, 33). Here, we highlight an alternative factor: selective “subunit” transformations that are applied to each input separately before they are combined. We chose non-invertible nonlinearities that exhibit generic selectivity (ON or OFF) and that, individually, induce lossy compression of stimuli with no inherent spatial statistical redundancy to exploit. Despite these properties, we found that a circuit with convergent, divergent architecture encoded more information with rectified subunit nonlinearities than with linear subunits.

This increase comes from a reformatting of the stimulus distribution in a manner that reduces the ambiguities produced by the convergence of multiple inputs (Fig. 2). In the LSC, it was possible to spread out the circuit responses by tuning the subunit weights (SI Appendix, Fig. S6) such that the ON subunits could be made independent of the OFF subunits. After applying the output nonlinearities, the response space for the LSC resembles that for the NSC. However, the entropy for the LSC still does not surpass that of the NSC (see SI Appendix, Figs. S6 and S7) because it does not reformat the mixed sign inputs as the NSC does (SI Appendix, Fig. S6). This reformatting facilitates the encoding of multiple stimulus features (mean luminance and contrast) in Figure 4. Thus, in the circuits we study here, efficient coding can be achieved with non-invertible nonlinear components. We note that even invertible nonlinearities, when followed by noise, will become difficult to invert and may thus behave like a non-invertible nonlinearity.

### Redundancy, correlation, and information

We find that the efficiency of divergent circuits can be enhanced by nonlinearities that decorrelate the outputs, as others have found (32, 38). Indeed, our findings show that diverging ON and OFF pathways resulted in efficiency gains for both the linear and nonlinear subunits circuits (compare the entropies for the single pathway configurations in Figure 2C to those for the corresponding divergent circuits in Figure 3F). Nonlinear responses in ganglion cells have more of an effect on decorrelating their responses than their center-surround receptive field properties (39). However, as pointed out in (39), weak correlation is not necessarily weak dependence. In the divergent, convergent circuits in Figure 3, rectifying nonlinearities located either in the output neurons or in the subunits decorrelate the outputs to a similar extent. However, a circuit with subunit nonlinearities produces the greater increase in entropy relative to a summation of linear subunits.

Maximizing information is often seen as equivalent to reducing redundancy (25, 28, 35, 40, 41). The responses from the NSC in Figure 3 have more information than those from the LSC and less redundancy (SI Appendix, Fig. S9). This is true despite their having the same degree of correlation, indicating that the reduction in redundancy is due to nonlinear reshaping of response distributions. The neural code in the retina is highly redundant (3, 35), as the degree to which neighboring ganglion cells share information has been estimated as roughly ten-fold (40). Our results suggest that the level of redundancy can be tuned by the subunit nonlinearities.

The connectivity structure and connection weights also have a role in reformatting inputs as they pass through a circuit. Compressed Sensing is a coding paradigm that has been used to model olfactory circuits in particular (42). In the presence of a compressive bottleneck in a neural circuit, Compressed Sensing is characterized by optimal connection weights that are sparse. Specifically, the highest levels of mutual information (or signal entropy) are obtained in these circuits when many of the weights potentially connecting inputs to neurons in the bottleneck are set to zero. Studies of Compressed Sensing with nonlinear units have related the parameters of such optimal sparse connectivity to observations and predictions in neural circuits (43, 44). One such study found that information was maximized by receptors that are uncorrelated and that selectively respond to half of the inputs (45). In SI Appendix Figure 8, we corroborate these findings and extend them to circuits with subunit nonlinearities. In a sparse, compressive circuit configuration, the inclusion of rectifying subunit nonlinearities leads to increases in encoded information relative to a sparse, compressive circuit with linear subunits. Here, we employed uniform weights with a wide range of sparsity levels, so as to highlight the contribution of the rectifying nonlinear subunits to the efficiency of the circuit responses in varied circuit architectures.

Bell and Sejnowski (19) showed that nonlinearities have the effect of reducing redundancy between output neurons by separating statistically independent parts of the inputs. Following that, it was shown that the efficient encoding of natural signals is facilitated by a nonlinear decomposition whose implementation is similar to the nonlinear behaviors observed in neural circuits through divisive normalization (46). Our study contributes to this body of work by showing how a circuit with convergent, divergent structure can leverage nonlinear subunits to contribute to a more informative, compressed representation by reducing redundancy (SI Appendix Fig. S9) independent of their effect on correlations.

### Reconciling selectivity with efficiency

Nonlinearities can have different functional consequences for neurons. Nonlinear transformations can induce selectivity in that they can cause a neuron to encode a very particular aspect of the stimulus or its inputs (47, 48). Nonlinearities can otherwise optimize efficiency by maximizing the entropy of the response distribution (23). The rectified nonlinearity that we used does not maximize the response entropy of the individual neuron that receives gaussian-distributed inputs, but it does enforce a strict selectivity for inputs above threshold. Selectivity would appear to be in conflict with efficient coding in that discarding information is a poor way to maximize it. Our results reveal how selectivity can work in concert with a circuit structure of parallel pathways to produce an efficient encoding of inputs.

The selective coding of features is often conflated with redundancy reduction, but it is important to make a distinction in the context of efficient coding - where a redundancy reducing code is reversible and is expected to maximize information about the stimulus (41). Selectivity indicates that some stimulus information will be irreversibly discarded. The existence of selective cell types that compute different aspects of the visual scene appears to confound an efficient coding framework (39). Yet, properties of selectivity are crucial to the functions of a diverse array of cell types, such as object-selective cells in medial temporal lobe (49), face-selective cells in the inferior temporal cortex (50, 51), and direction-selective cells, orientation-selective cells, and edge detector cells in the retina (52). Furthermore, many cell types in the retina and other circuits have both an ON and an OFF variant, indicating that this kind of ON/OFF selectivity is beneficial to sensory information processing (20, 32).

### Implications for artificial neural networks

Although mean and contrast are elementary features of visual inputs, the striations seen in the response space in Figure 4 (NSC, right plots) reflect the concept that hidden nonlinear neural units can facilitate the categorization of stimulus features (53). In our study simply inserting nonlinear subunits with uniform weights immediately produced a representation that may enable linear classification or decoding of the mean and contrast levels of the input.

Feedforward artificial neural networks (ANNs) were inspired by the layered organization of biological neural networks. Neural units have activation functions, or static nonlinearities, that transform inputs. Rectified Linear Units (ReLU) such as those used in our nonlinear neural units, enforce a strict selectivity for inputs above threshold; whereas smooth nonlinearities implement a less rigid selectivity, if at all. In both cases, selectivity is dependent on the bias and weight parameters, which can be adjusted by learning, to offset the nonlinearity such that it truncates the input distribution to various degrees. The ReLU frequently has the best performance among other nonlinear activation functions (54, 55) in tasks ranging from the discrimination of handwritten digits to restricted Boltzmann machines (56). The findings presented here of the information preserving capabilities of a selectivity-inducing nonlinear activation within an architecture that is reminiscent of a feedforward ANN complement our knowledge of the ReLU’s favorable performance in machine learning and the remarkable classification capabilities of ANNs.

### Future directions

The interaction between noise and the nonlinearities, convergence, and divergence studied here is potentially very interesting. Our results did not depend on noise explicitly; however, we note that the discretization of the response distributions effectuates a low level of output noise because stimuli that fall into the same discrete bin cannot be disambiguated. In SI Appendix Figure S1, we explicitly introduce weak noise after the subunit summation and confirm that our main results continue hold: the NSC maintains an advantage over the LSC.

Overall, the magnitude and source of noise can have a large effect on a circuit’s ability to encode stimulus information. As a preliminary check, we confirmed that one of these known effects carries over to our convergent/divergent circuit. In a theoretical study of divergent ON/OFF neuron motifs, Brinkman et al (37) found that for low noise conditions, mutual information is optimized by nonlinearities that cross at their “lower bend,” similar to the default crossing at zero threshold for the rectifying ON and OFF nonlinearities in our study. For high noise conditions, the mutual information is optimized by nonlinearities that overlap, suggesting redundancy in these cases. We confirmed this effect in a convergent/divergent circuit with noise after the summed subunits (SI Appendix, Fig. S10). Our future studies will build on these preliminary explorations to more completely describe the effects of noise on optimal coding within these circuits.

Additionally, our model did not include temporal dynamics. We opted for a granular, geometrical analysis of the set of all possible responses to a fixed and finite set of stimuli so that we could clearly ascertain the counterintuitive finding that a convergent, divergent circuit can preserve more information with rectified nonlinear subunits than with linear subunits. Despite the lack of temporal dynamics, we compared the effects of different output nonlinearities which abstractly approximate different spike generating mechanisms. Future studies will explicitly include time-dependence to investigate how adapting subunit nonlinearities impact the efficient encoding of inputs with changing stimulus statistics.

## Materials and Methods

We used Shannon’s information (16) to quantify the information retention of our model circuits because it quantifies how many distinct neural responses are possible given a particular stimulus distribution, and this relates to the specificity of encoding even though it does not indicate which specific stimulus features are encoded. Since there was no noise anywhere in the circuit, the mutual information between the stimulus and the response reduces to the entropy of the response. Mutual information is defined as (17):

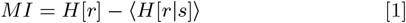

Where *r* is the output response and *s* is the stimulus input. *H*[*r*] is the response entropy and *H*[*r*|*s*] is the conditional entropy of the response given a stimulus, *s*.

In our study, there is a deterministic relationship between the response and the stimulus due to the lack of noise. The second term of the MI goes to zero and one is left with the entropy of the response. We were careful to avoid any effects that could distort the interpretation of the entropy. The ReLU was ideal here because it compresses the input signal without necessarily scaling it. In our model, the compression was entirely derived from the non-invertibility of the nonlinearity rather than a linear gain factor.

The convergent structure of the retina reduces the dimension of the high-resolution visual input it receives, placing an upper bound on the amount of information that can possibly be transmitted through the optic nerve. In general, the data compression implemented by the circuit architecture may perform lossless or lossy compression or some combination, depending on the statistics of the inputs. In this study, we focus on lossy compression. By using sample “images” of uncorrelated gaussian random inputs (i.e. no redundant structure), we place the inputs into a regime where lossless compression is impossible or assumed to have already taken place. Therefore, the circuit configuration that experiences less information loss has a higher entropy than that which experiences more information loss relative to the information contained in the stimulus. We thus consider higher entropy to be an indication of better performance.

### Model simulations and visualizations

All simulations, visualizations, and entropy computations were done in Matlab. The dimension of the stimulus always matches the dimension of the subunits within a pathway, and a stimulus consists of N stimulus inputs. For example, if there are 5 subunits in each of the ON and OFF pathways, then the stimulus has 5 stimulus inputs (sometimes referred to as pixels). Each stimulus input was independently drawn from a gaussian distribution with arbitrary units (*µ* = 0, *σ* = 10). Each subunit receives input from one stimulus input. For all figures in this paper, linear subunits did not transform stimulus inputs and therefore the ON linear subunit response was equivalent to the stimulus input and the OFF linear subunit response was the negative of the stimulus input.

All weights were uniform with unit weights from stimulus inputs to subunits and normalized weights from subunits to outputs. The subunits were normalized so that the variance of the linear sum of subunits is maintained. With *N* subunits, each subunit weight is 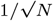. This normalization facilitated a comparison between circuit configurations with linear and nonlinear subunits and varying numbers of convergent subunits. All circuit configurations are subject to the same uniform weighting and subunit normalization throughout the paper except where noted in the Supplemental Information.

Each rectified nonlinear unit has unit slope (slope = ±1) and applies a threshold to the stimulus input - effectively a positive-pass filter for ON subunits and a negative-pass filter for OFF subunits.

The output neuron linearly sums the subunit responses in its pathway and then applies the output nonlinearity. The output response to a given stimulus is a single value that represents a steady state response, as our model does not have temporal dynamics.

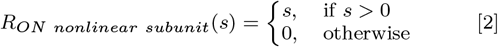

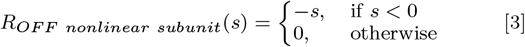

For the summed subunit response before or without an output nonlinearity in a single pathway,

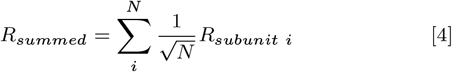

And with an output nonlinearity, the single pathway output response is

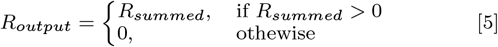

The optimal sigmoidal output nonlinearity was computed as the cumulative gaussian of the summed subunit distribution, bounded by the maximum and minimum summed subunit values:

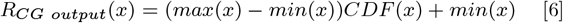

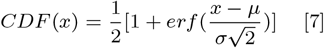

### Visualizations in stimulus, subunit, and response spaces

Each quadrant was color-coded such that: *s*_1_ *>* 0, *s*_2_ *>* 0: blue; *s*_1_ *<* 0, *s*_2_ *>* 0: orange; *s*_1_ *<* 0, *s*_2_ *<* 0: yellow; *s*_1_ *>* 0, *s*_2_ *<* 0: purple. Output response histograms in Figure 3 are also color-coded in this way to show which response bins represent which stimuli. For mean luminance and contrast visualization, spaces were color-coded to indicate bands of mean stimulus luminances, *M*, and stimulus contrasts, Λ. Each stimulus image, γ, consists of *N* stimulus inputs, γ = [*s*_1_, *s*_2_, …, *s*_*N*_]. In Figures 3A-E and 4, *N* = 2.

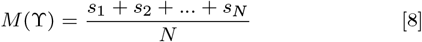

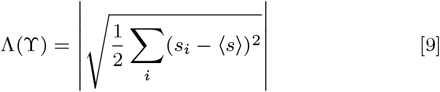

### Entropy calculations

Entropy computations were done by simulating the circuit responses to batches of stimulus samples. Discrete entropy was used to quantify continuous stimulus and response distributions. Distributions of stimuli and responses were binned and probabilities were computed from the binned distributions. These binned probability distributions were used to calculate the entropy of the responses. Information entropy is defined as

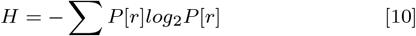

where P[r] is a discrete probability distribution.

The entropy quantities presented are the average over 10 batches of samples. The normalized entropy was computed by dividing the entropy of the circuit by the entropy of the sum of linear subunits. Thus,

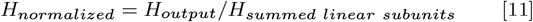

The entropy of the summed linear subunits was the same for the single pathway as it was for the divergent pathways (*H*_*summed linear subunits*_ = 7.37 bits) since the OFF linear subunits are perfectly anti-correlated with the ON linear subunits and do not provide additional information. The subunit normalization facilitated a comparison between the entropies of the circuit configurations in Figures 2 and 3 and across different quantities of converging inputs.

A Freedman-Diaconis histogram bin approximation was used to determine an appropriate bin width for the stimulus and response distributions (57). A consistent bin width of 0.25 was used for all entropy calculations to facilitate comparison. This bin width was used for all dimensions. For example, in a 2-dimensional response space, bins would be boxes that are 0.25 × 0.25. These discretization parameters were chosen carefully to ensure that the bins were sufficiently small to capture the shape of the distributions, but not so small that the log(N) bound was reached. The Freedman-Diaconis estimation returns a bin width that minimizes the difference between the areas under the curves of the discrete and continuous distributions.

To ensure confidence in the entropy calculation, the sample batch size was computed as follows. First, binned entropies were computed for gaussian distributions with a range of variances and a range of batch sizes. Then, the entropy error was computed as the absolute difference between these numerical binned entropies and their corresponding analytic binned entropies (eqn. 12). Linear fits of the entropy error as a function of batch size were computed for each value of distribution variance individually. Then another linear fit was performed for those first fit parameters as a function of distribution variance. This procedure produced a general expression for the entropy error given distribution variance and batch size. We chose an entropy error tolerance of 0.005 which we used to determine an appropriate batch size. The minimum batch size for entropy computations in the main text was 10_6_ samples. Smaller batch sizes were permitted for noise entropy computations in the Supplemental Appendix.

The 36-dimensional input space used in Figure 1E,F was too large for a numerical computation of the entropy. Its discrete entropy was estimated analytically from its continuous entropy with a bin-correction term as in equation 12 where m = 36, bin width b = 0.25, and K is the covariance matrix of the stimuli.

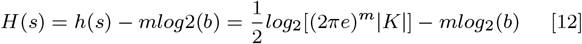

### Data Availability

All simulations and analyses were done in Matlab using custom-written scripts. These can be found on the corresponding author’s Github page: https://github.com/gabrielle9/nonlinear-convergence-info-entropy-retention.

## ACKNOWLEDGMENTS

We thank Joel Zylberberg, Stephen Baccus, Max Turner, Adree Songco-Aguas, Iris Jianghong Shi, and Matthew Farrell for their helpful feedback and comments on the manuscript and Leenoy Meshulam for their helpful discussions.

Funding sources: GJG is supported by NIH NINDS K22 1K22NS104187-01A1 and WRF UWIN postdoc fellowship, ETSB was supported by the Boeing professorship in Applied Mathematics at UW and the UW Center for Sensorimotor Neural Engineering. FR is supported by EY028111.

## Supplementary Information for

## Appendix I: Analytic derivation of variance of summed nonlinear subunits distribution

The nonlinear subunit responses are described by a rectified gaussian distribution which has variance, 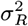.

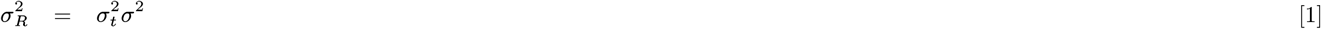

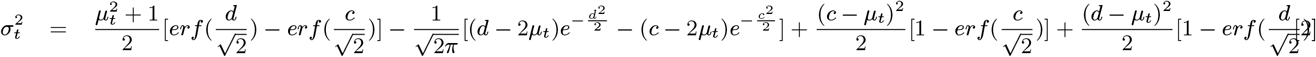

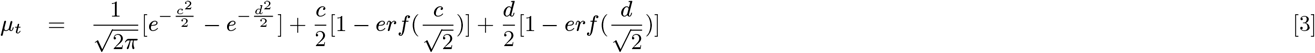

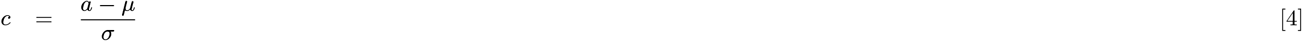

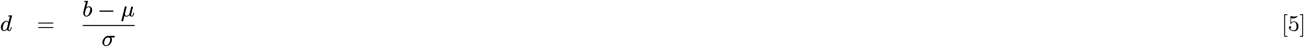

where *σ*^2^ and *µ* are the variance and mean, respectively, of the unrectified distribution which would be the same as the variance and mean of the summed linear subunits distribution. The lower and upper bounds of the rectified gaussian distribution are given by *a* and *b*, respectively, however, in our case the upper bound is infinite.

With *µ* = 0, *σ* = 10, *a* = 0, *b* = *∞*, we obtain, *c* = 0, *d* = *∞*, 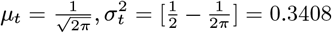. Thus, 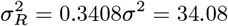.

## Appendix II: Analytic arguments for higher entropy in nonlinear subunits circuit

The circuit with divergent pathways and nonlinear subunits was shown to have greater entropy than the circuit with linear subunits in the numerical computations in the main paper. We provide supporting analytic arguments here, in the approximation that one takes the limit of a large (infinite) number of normalized subunits. By the central limit theorem, the summed output of the nonlinear subunits then approaches a gaussian distribution. The general continuous entropy expression for a gaussian distribution is:

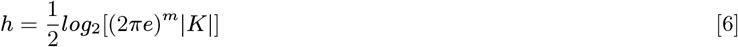

where *m* is the dimension of the gaussian distribution. With ON and OFF outputs, *m* = 2. *K* is the determinant of the covariance matrix of the output distribution, with variance *σ*^2^ and correlation coefficient *ρ* between the ON and OFF outputs (0 *≤ ρ*^2^ *≤* 1):

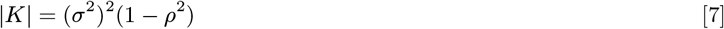

For the sum of linear subunits, *ρ*_*lin*_ = *–*1, and *σ*^2^ = 100. For the sum of nonlinear subunits, we denote 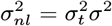 (see Appendix I for analytical derivation of the factor 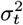). These values hold for all subunit dimensions because the subunits are normalized.

The entropy equation can be expanded to:

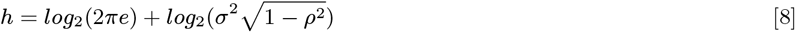

The condition that the entropy of the nonlinear subunits circuit is higher than that of the linear subunits circuit is equivalent to this inequality:

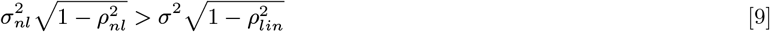

Which can be rearranged:

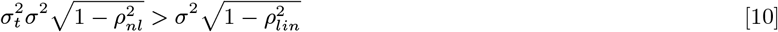

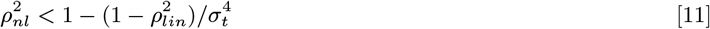

This condition is immediately satisfied since 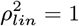. The final step would be to apply the output nonlinearities, however, this would not alter the sum of nonlinear subunits and would only serve to reduce the entropy of the sum of linear subunits. Therefore the sum of nonlinear subunits will always have a higher entropy than the sum of linear subunits (for the particular circuit configurations and noiseless conditions studied here).

To extend this analysis, we determine how much decorrelation of the ON and OFF linear pathways is tolerated before the entropy of the summed linear subunits overcomes the entropy of the summed nonlinear subunits. By rearranging equation 11 and substituting in the values of 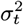 (from Appendix I) and the numerically computed value of *ρ*_*nl*_ (0.4670, see Results in main text), we arrive at the following condition:

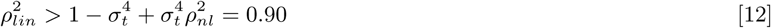

Therefore, the linear pathways can tolerate about 10% decorrelation before overtaking the entropy of the nonlinear subunits circuit. We remind the reader that this is the case when there are no output nonlinearities. As soon as the output nonlinearities are taken into account, the linear subunits circuit remains lower than that of the nonlinear subunits circuit regardless of the extent of decorrelation induced by the orthogonalization of the weights (see Supplemental Fig. 5 and 6).

## Supporting Figures

**Fig. S1.**
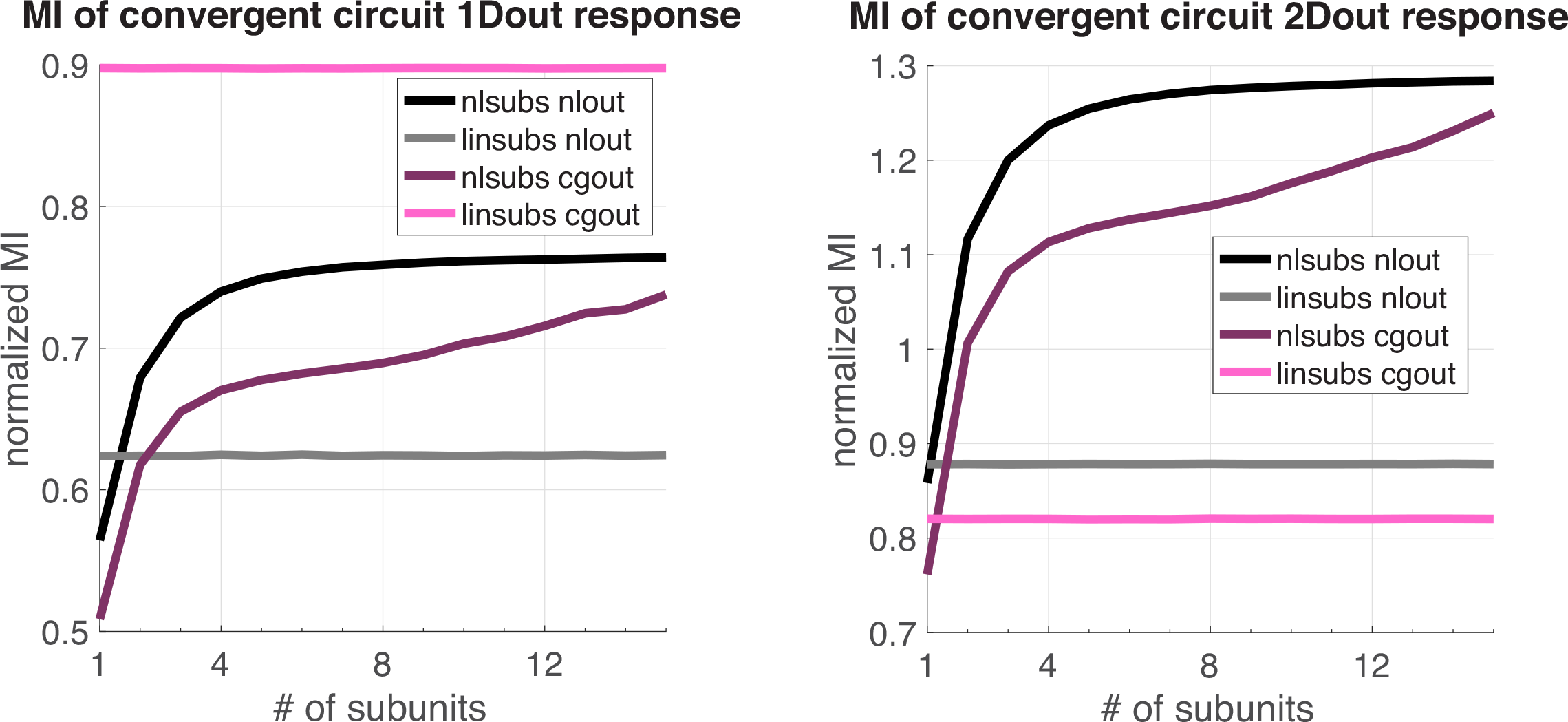
Mutual Information for different circuit configurations with noisy outputs. Left: single pathway circuit; Right: ON and OFF pathway circuit. Black: rectifying nonlinear subunits and rectifying nonlinear output; Grey: linear subunits and rectifying nonlinear output; Dark magenta: rectifying nonlinear subunits and cumulative gaussian output nonlinearity; Pink: linear subunits and cumulative gaussian output nonlinearity. Stimulus distribution has *σ*_*s*_ = 10, noise distribution has *σ*_*m*_ = 1. Noise arrives after the subunit summation but before the output nonlinearity. The cumulative gaussian output nonlinearities are the same as in Figures 2 and 3 in the main text and were optimized for the output response distribution alone. As a result, the cumulative gaussian nonlinearity is not optimal for the response distributions that also contain noise. As with the normalized entropies for the different circuit configurations in Figures 2 and 3 in the main text, the NSC encodes more information as the number of convergent subunits increases and it encodes more information than the LSC. There is an exception when there is only a single subunit due to the placement of the noise after the subunit response but before the output nonlinearity is applied.

**Fig. S2.**
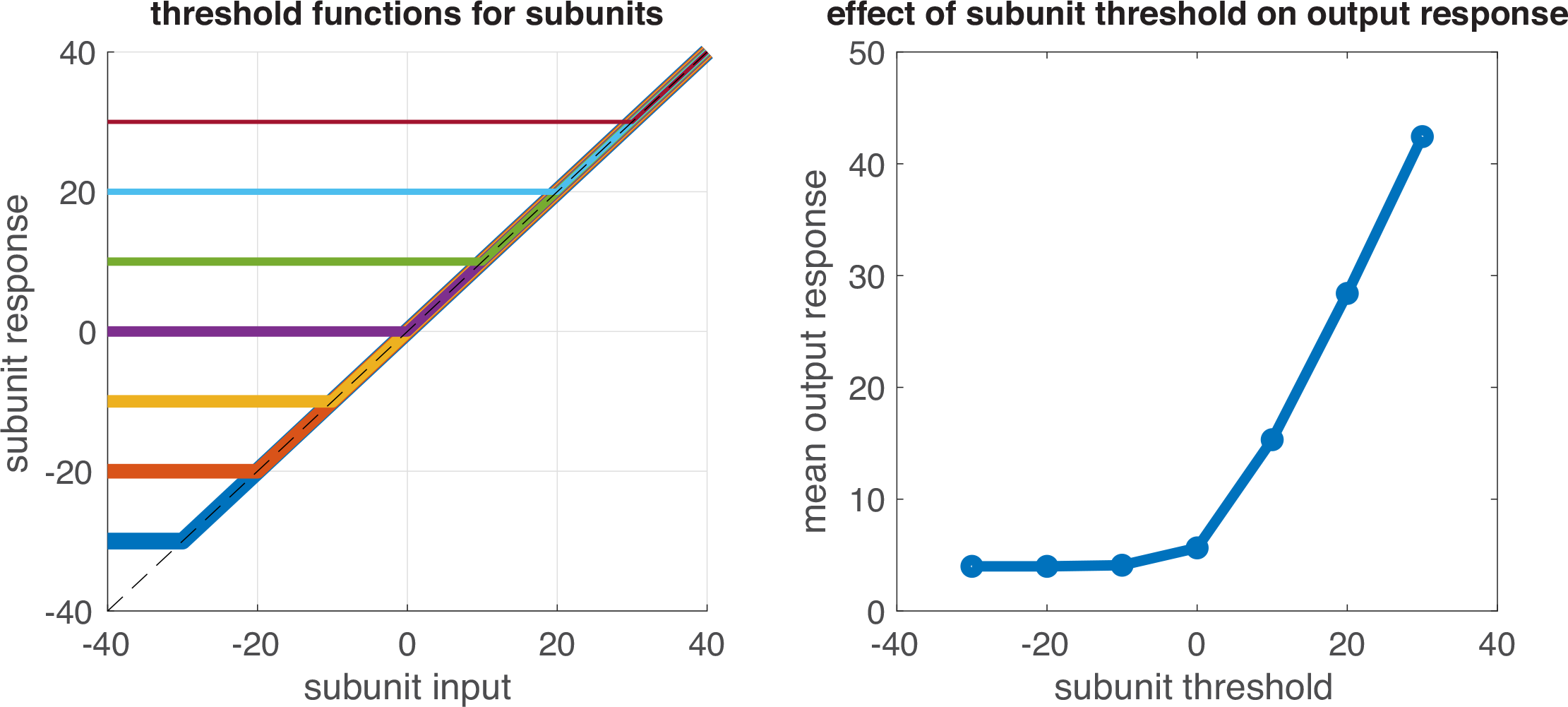
Implementation of sliding subunit thresholds in reference to Figures S2, S3, and S4. Left, threshold implementation for subunits showing how low subunit thresholds resemble a linear response function whereas high subunit thresholds rectify much of the subunit input range. Right, mean output response of ON output neuron as a function of subunit threshold (negative thresholds indicate subunits are more linear). There are 2 ON subunits converging to a single output neuron with a rectifying nonlinearity. The output nonlinearity threshold is fixed at zero. Output response is in arbitrary rate units. As the subunit threshold increases from linear to highly rectifying, the output neuron activity increases nonlinearly. Simulations for each subunit threshold value were run using a 2D gaussian distribution of stimuli as the stimulus inputs. The mean output response for the ON output neuron was computed by taking the mean of the ON output response distribution.

**Fig. S3.**
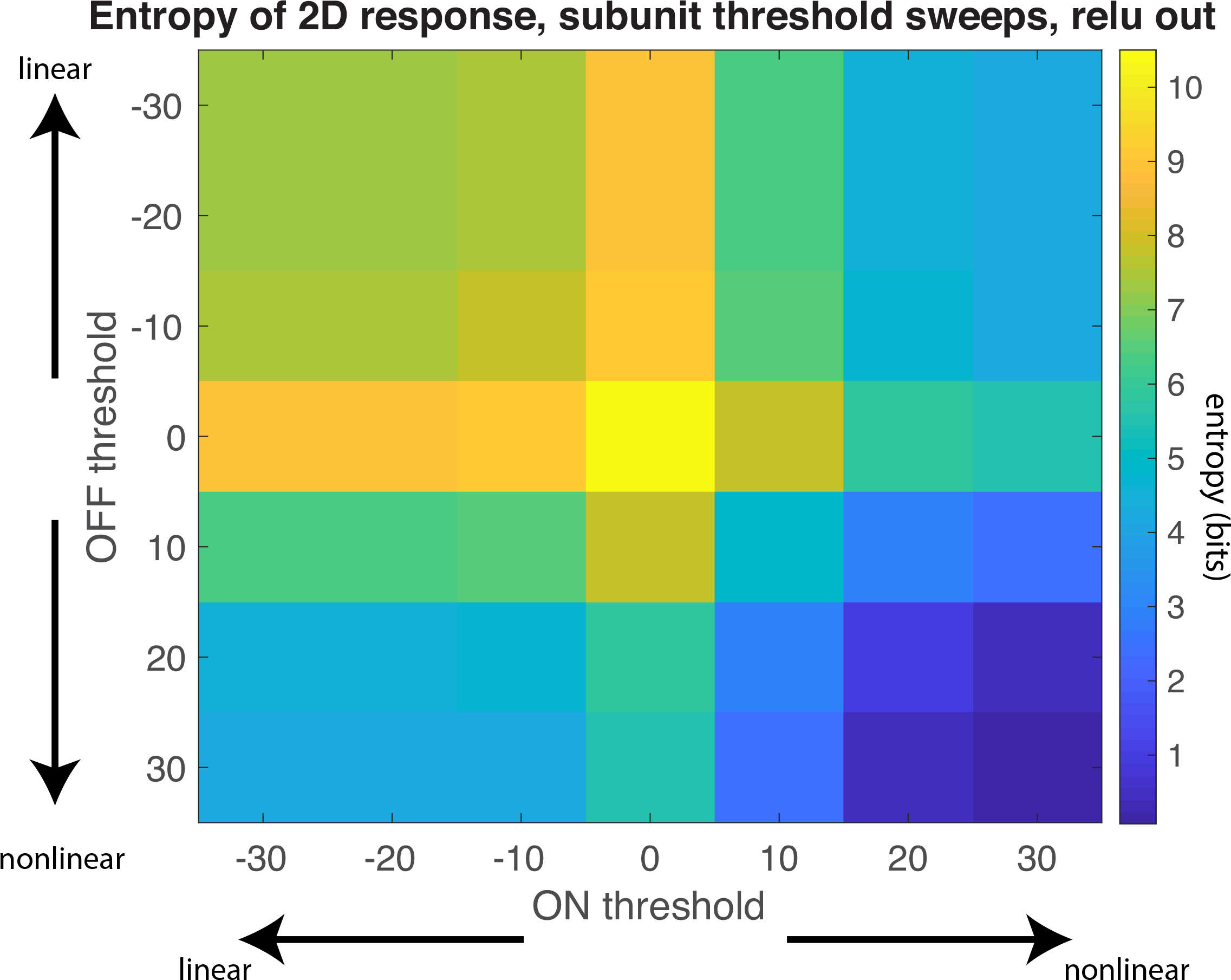
Entropy of circuit response for different nonlinear subunit thresholds ranging from -30 to 30 (arbitrary response units). Circuit has 2 inputs, an ON and an OFF pathway, and fixed output nonlinearities thresholded at zero for each pathway. Negative thresholds approach linear subunits while positive thresholds are extremely rectified. All subunits within the same pathway have the same threshold but ON subunit thresholds can vary independently from OFF subunit thresholds in these sweeps. The highest circuit response entropy is produced when ON and OFF nonlinear subunit thresholds are at zero, which is the mean of the input distribution.

**Fig. S4.**
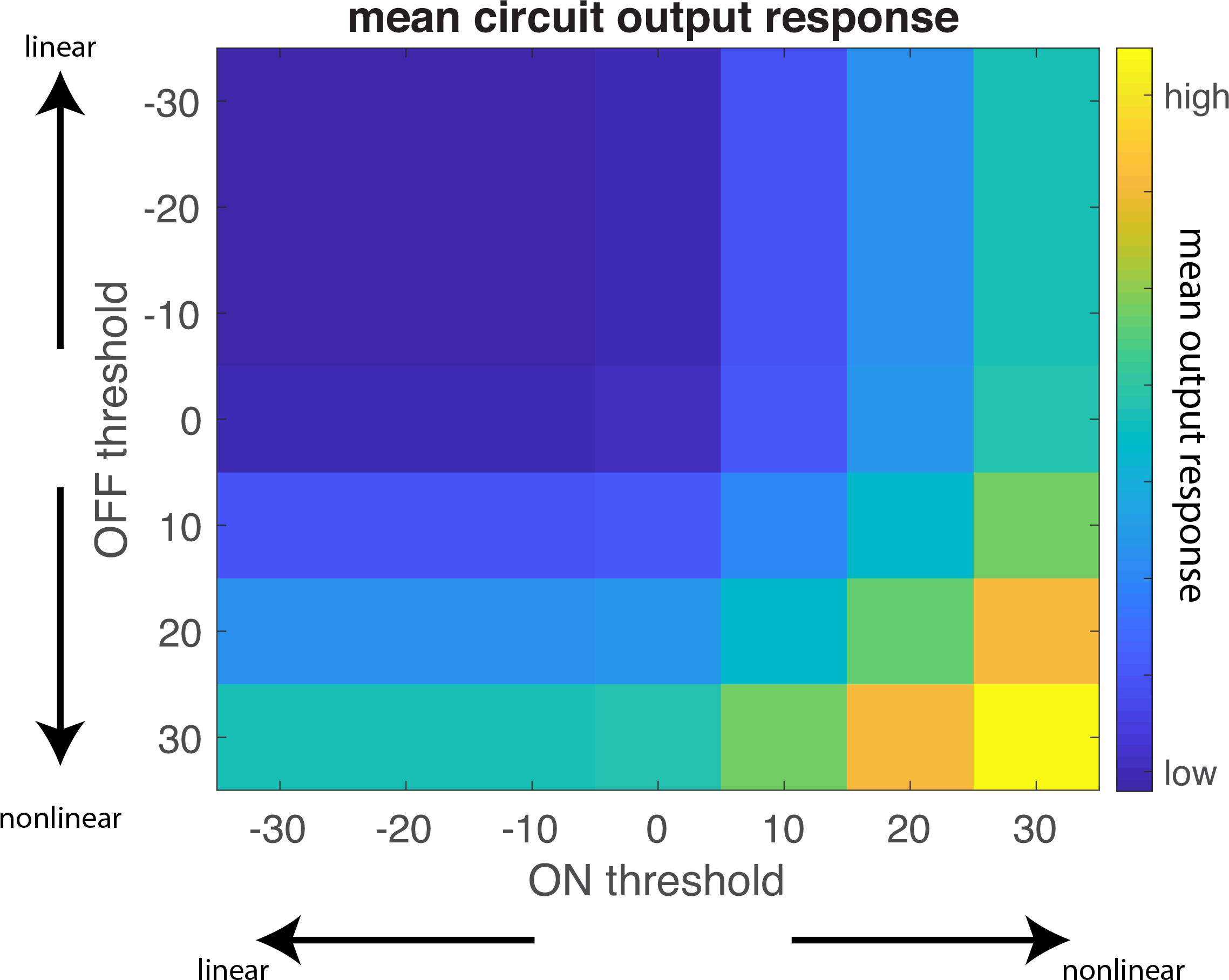
Mean circuit output response as a function of ON and OFF subunit thresholds. Circuit has 2 inputs, an ON and an OFF pathway, and fixed output nonlinearities thresholded at zero for each pathway. Mean output response is displayed in arbitrary response units on the colorbar. Mean output response is computed as mean[ON output, OFF output]. Simulations for each combination of subunit thresholds were run using a 2D gaussian distribution of stimuli as the stimulus inputs. The mean of the ON output response distribution and the OFF output response distribution was computed before taking the mean among the ON and OFF outputs. Negative thresholds indicate more linear subunits while positive thresholds indicate more extreme subunit rectification. The extremely rectified subunits produce higher mean output responses; whereas moderately rectified or fully linear subunits produce relatively low mean output responses.

**Fig. S5.**
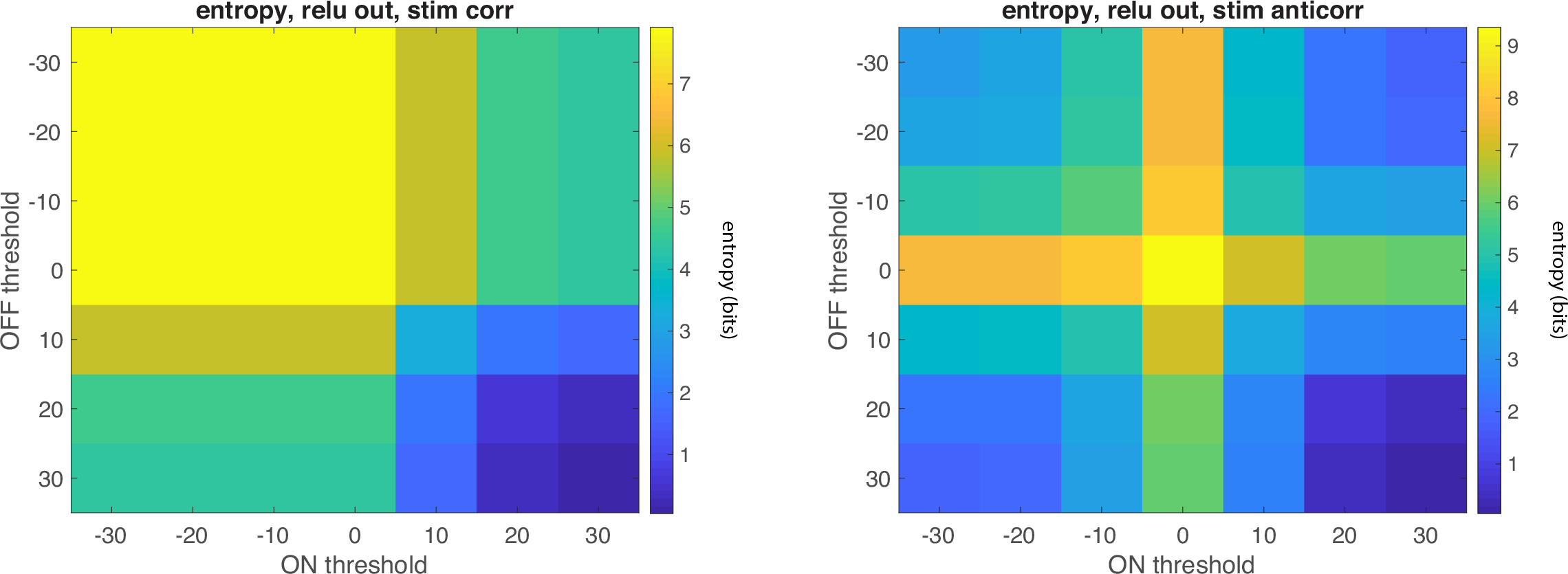
Entropy as a function of subunit thresholds for ON and OFF pathways in a circuit with 2 inputs, an ON and an OFF pathway, and fixed output nonlinearities thresholded at zero for each pathway. Left, stimuli are highly correlated (cc = 0.995). Right, stimuli are highly anti-correlated (cc = -0.995). Colorbars show discrete entropy values in bits. For correlated stimuli, the entropy of the circuit is not very sensitive to the linearity of the subunits except when subunits are extremely rectifying. This reinforces the observation that nonlinear subunits thresholded at zero and linear subunits encode correlated stimuli similarly (see Fig. 3 in main text). In contrast, the circuit entropy is sensitive to anti-correlated stimuli and the entropy is highest when the ON and OFF subunit thresholds cross at zero.

**Fig. S6.**
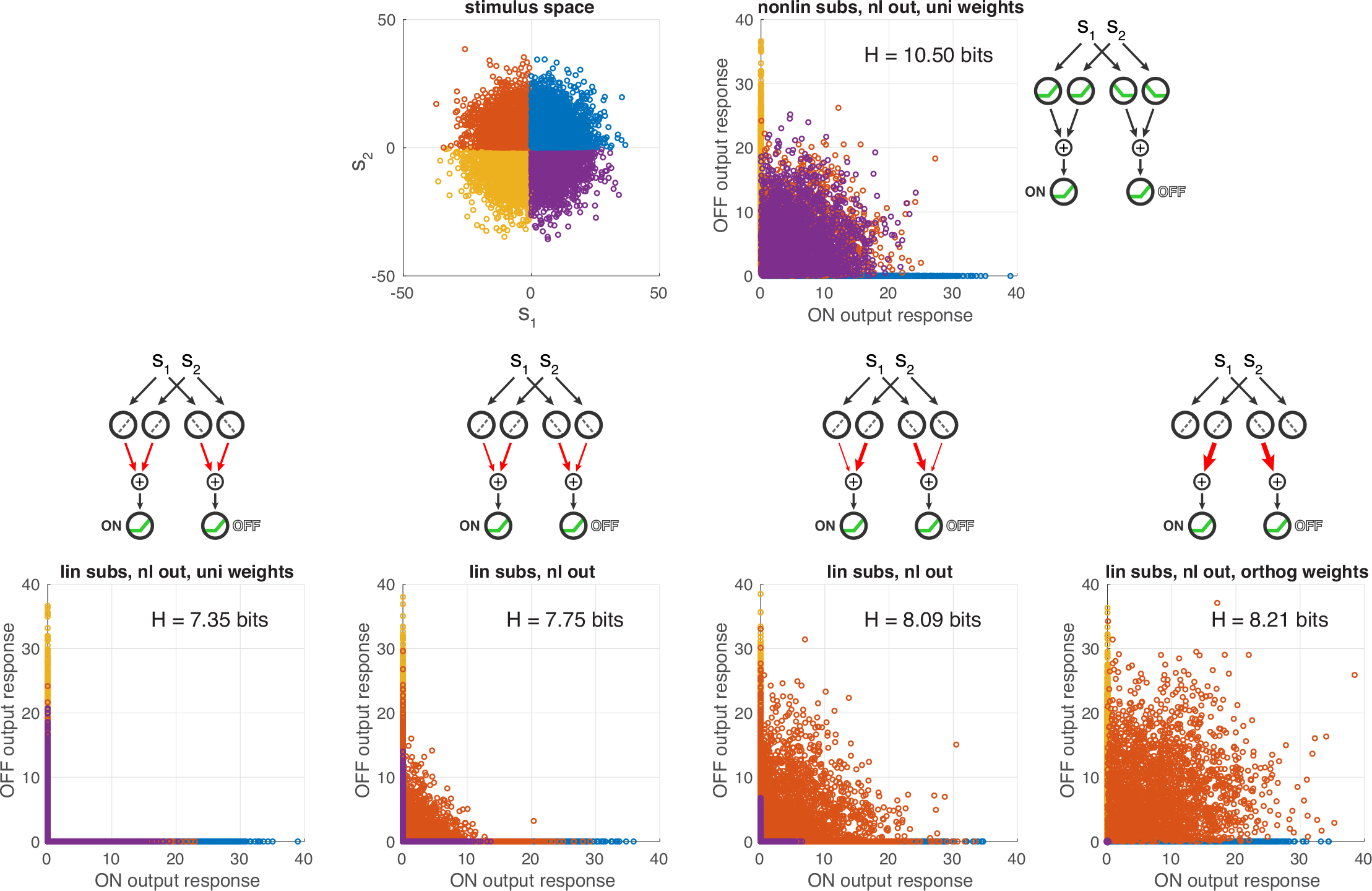
The linear subunits can be reweighted to form an output distribution that resembles that for the NSC. Top left, stimulus space with color-coded quadrants; top right, output response space for NSC with 2 input dimensions and uniform subunit weights (as shown in schematic; also see Fig. 3E in main text). Bottom, output response space for LSC with 2 input dimensions and (left) uniform subunit weights, (2nd and 3rd from left) subunit weights rotating away from uniform, (right) orthogonal subunit weights (see Fig. S7 for depiction of weights rotation). Despite the resemblance between the response spaces in the top right and bottom right, orthogonalizing the linear subunit weights still produces lower entropy than the NSC with uniform weights. The color coding reveals that for the LSC, as the orange points are liberated from the axes, the purple points are compressed to the origin, in contrast to the case for the NSC where both the orange and purple points are pushed away from the axes. As schematized, all circuits have 2 inputs, an ON and an OFF pathway, and fixed output nonlinearities thresholded at zero.

**Fig. S7.**
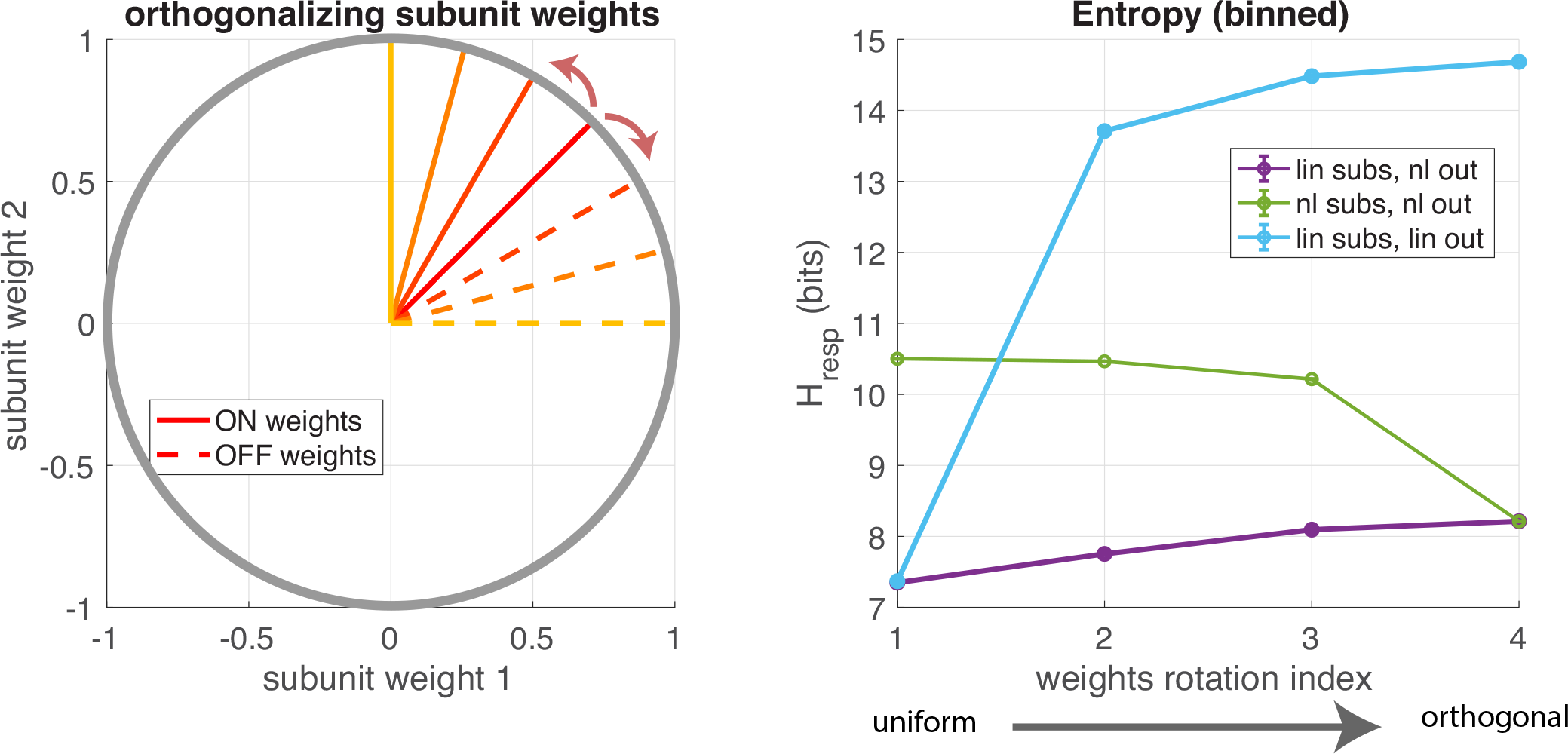
Entropy for 3 circuit configurations as subunit weights are rotated from uniform to orthogonal. All circuit configurations have 2 inputs, and an ON and an OFF pathway. Left, since subunits are normalized by 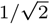 they are bounded by the unit circle. The uniform subunit weights are at 45 degrees whereas the orthogonal subunit weights are at 0 and 90 degrees. More explicitly, the uniform weights have 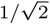 for all subunit weights while the orthogonal weights have [0,1] for the ON subunits and [1,0] for the OFF subunits (see schematic in bottom right panel of Fig. S6 for a depiction of orthogonal weights in the circuit). Right, discrete entropy as a function of subunit weight orientation. The weights rotation index begins at the uniform subunit weights and ends at the orthogonal subunit weights. The LSC (purple curve) maintains the lowest entropy among the circuit configurations, consistent with Supplemental Figure S6. The entropy for the NSC (green curve) drops to meet the LSC when the subunit weights are completely orthogonal. In reference to the derivation in Appendix II, the entropy of the fully linear circuit (linear subunits and linear output) is shown in blue. As the subunit weights are rotated, the entropy quickly increases because there is no output nonlinearity to constraint the output space as the ON and OFF pathways become entirely independent.

**Fig. S8.**
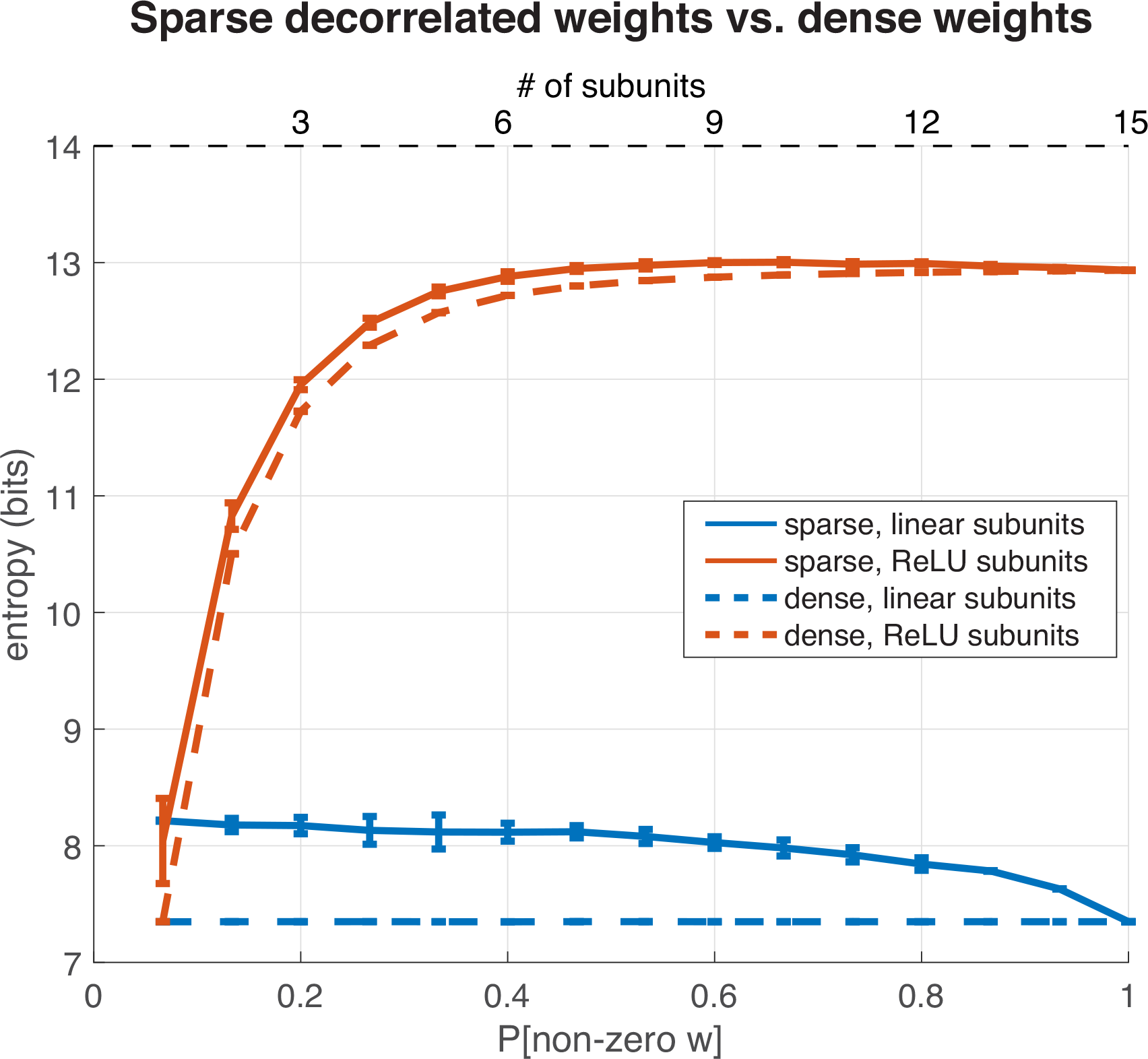
Entropy for circuit configurations with linear and nonlinear subunits. These circuits have an ON and an OFF pathway and rectifying nonlinear outputs. The dashed curves are the non-normalized versions of the black and grey curves from Figure 3F in the main text and they correspond to the top x-axis which indicates the number of convergent subunits. These circuits have the same uniform weightings that were used throughout the main text of the paper. The solid curves represent a sparse weighting of the subunits that decorrelates the ON and OFF pathways. There, the circuits have 15 subunits but the lower x-axis indicates the proportion of those 15 subunit weights that will be non-zero. At P[non-zero w] = 1, the weights are fully dense (matching the cases with 15 subunits in the main paper and in the dashed curves), but for lower P[non-zero w] the weights are sparser. The subunits with zero weights are randomly chosen and they are independent between the ON and OFF pathways. Each point is the average of 10 simulations. Error bars represent the standard deviation among the 10 simulations. Input signal has *σ*_*s*_ = 10, no circuit noise. This figure compares the entropy of the divergent circuit when the ON and OFF pathways receive correlated inputs (dense weights) to the entropy when the ON and OFF pathways receive decorrelated inputs (sparse weights). More specifically, it allows one to see how the convergence of some number of subunits is impacted by the correlations, or lack thereof, between the ON and OFF subunits. Starting at the right side of the plot, as the number of subunits is decreased in the dense weight circuits, the entropy decreases for the NSC (as it does in Fig. 3F in the main text). Meanwhile, as P[non-zero w] decreases and weights become sparser for the sparse circuits, there is an increase in both the LSC and the NSC entropy relative to the entropy of the dense circuits. However, the sparse LSC entropy does not increase enough to meet that of the NSC until the lowest P[non-zero w] is reached - which is where the two circuits are equivalent because the convergence step cannot differentiate them since there is only 1 non-zero subunit.

**Fig. S9.**
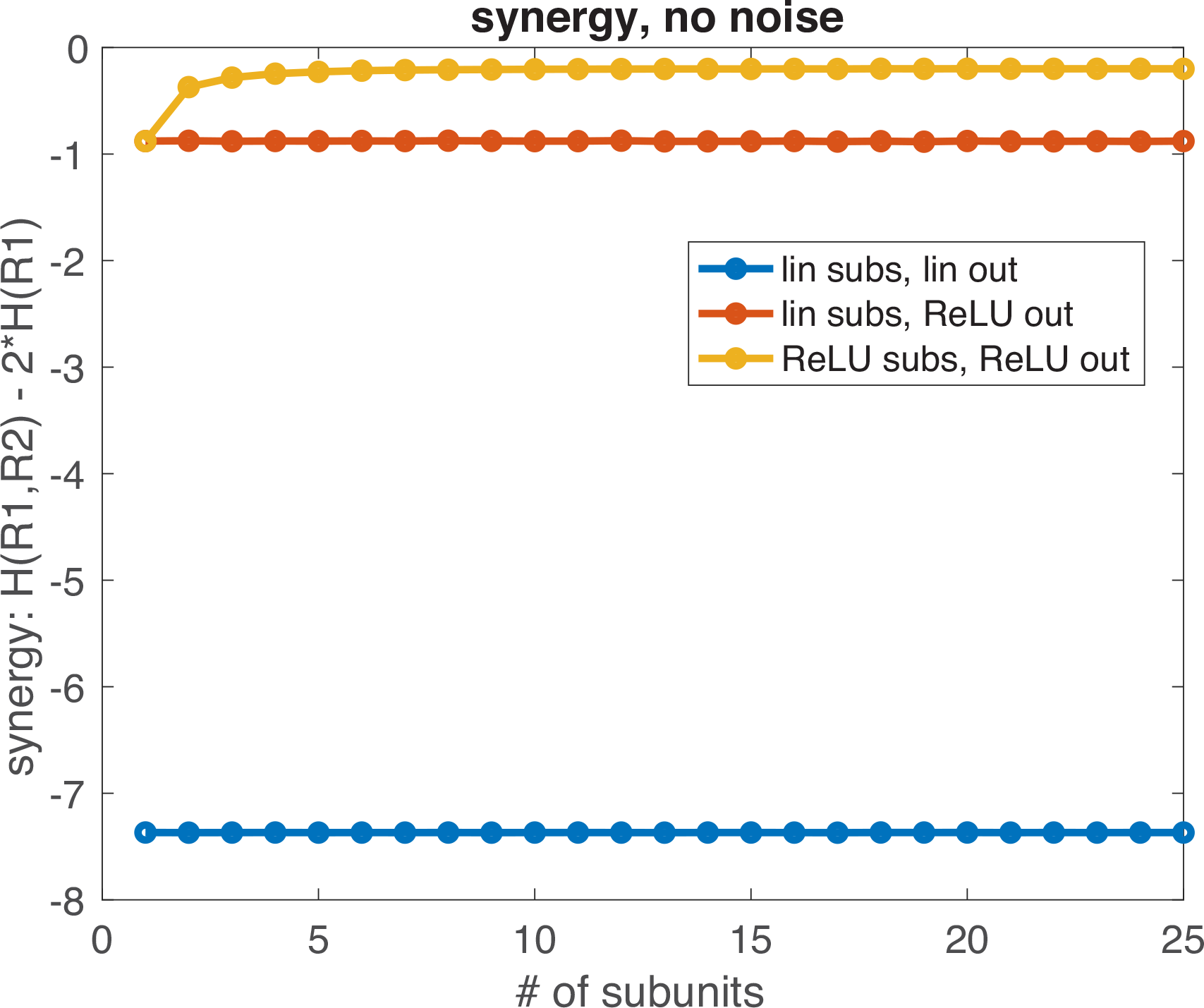
The synergy for 3 circuit configurations as a function of the number of convergent subunits. All circuits have 2 inputs, and diverging ON and OFF pathways. Synergy(R1,R2) = I(S;R1,R2) - I(S;R1) - I(S;R2) Where I stands for mutual information, R is the output response, and H is entropy. The output responses are deterministic and thus the synergy reduces to: Synergy(R1,R2) = H(R1,R2) - H(R1) - H(R2) Positive synergy values would indicate that there is more information in the ON and OFF outputs jointly than the sum of the information computed in the ON and OFF outputs separately. Negative synergy values indicate redundancy among the ON and OFF outputs (1–4). The fully linear circuit (blue) has the most redundancy because the ON and OFF outputs contain the same information and are simply anti-correlated. With linear subunits and a rectified output nonlinearity (orange), the redundancy is greatly reduced - it would be zero if it were not for the overlap in outputs for the stimuli that sum to zero. The NSC (yellow) has increasing redundancy as the number of subunits increases. As more responses are freed from the output response manifold, the independence between the ON and OFF outputs saturates.

**Fig. S10.**
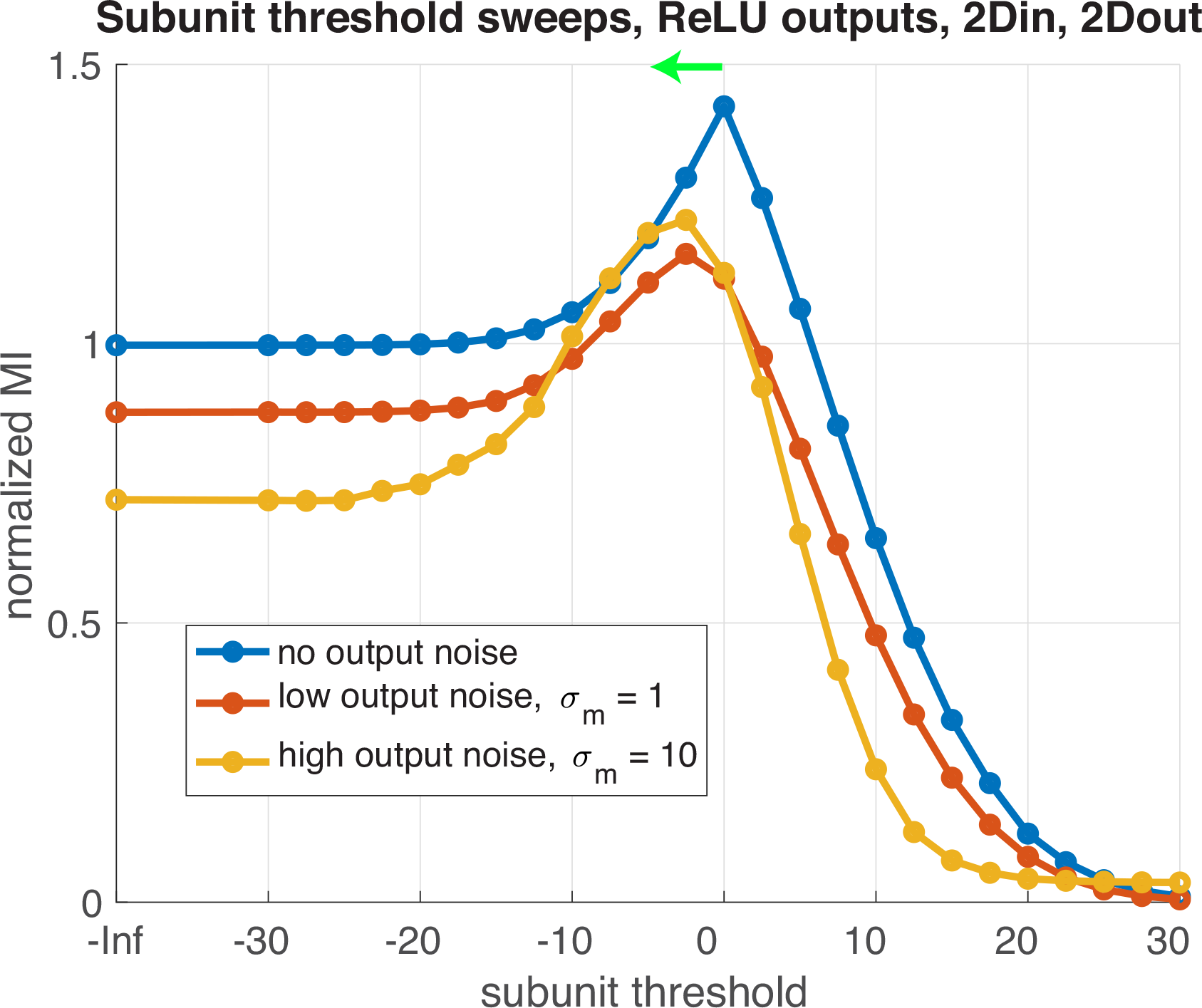
Optimal threshold is more linear with higher noise. In a circuit with 2 nonlinear outputs and 2 subunit inputs each, the optimal subunit threshold depends on the amount of noise after the subunit summation. As in SI Fig. S1, output noise is applied after the subunit summation but before the output nonlinearity is applied. With no noise, the optimal subunit threshold is zero, as corroborated by SI Fig. S3. As noise is increased, the optimal subunit threshold shifts lower towards more linear subunits. A lower subunit threshold allows the ON and OFF pathways to encode some overlapping information which may help the circuit to retain more information when noise has a corrupting effect on the input signals by introducing redundancy.

